# Assessing post-stroke cognition in pre-clinical models: lessons and recommendations from a multi-center study

**DOI:** 10.1101/2025.05.21.655267

**Authors:** G. Brezzo, K. A. Zera, D. Straus, J. E. Goertz, S. H. Loppi, R. Crumpacker, J. B. Frye, D. A. Becktel, M. I. Cuartero, A. García Culebras, C. Dames, D. Berchtold, J. H. Fowler, A. Meisel, J. Anrather, M. A. Moro, S. M. Allan, K. P. Doyle, M.S. Buckwalter, B.W. McColl

**Affiliations:** UK Dementia Research Institute, University of Edinburgh, Edinburgh EH16 4SB, UK; Centre for Discovery Brain Sciences, University of Edinburgh, Edinburgh EH16 4SB, UK; Department of Neurology and Neurological Sciences, Stanford University School of Medicine, Stanford, CA, USA; The Feil Family Brain and Mind Research Institute, Weill Cornell Medicine, New York, NY 10065 USA; Department of Immunobiology, University of Arizona, Tucson, AZ, USA; Neurovascular Pathophysiology, Cardiovascular Risk Factors and Brain Function Programme, Centro Nacional de Investigaciones Cardiovasculares (CNIC), Calle Melchor Fernández Almagro, 3, 28029 Madrid, Spain; Unidad de Investigación Neurovascular, Departamento de Farmacología, Facultad de Medicina, Universidad Complutense de Madrid (UCM), Plaza Ramón y Cajal, s/n, Moncloa - Aravaca, 28040 Madrid, Spain; Departamento de Biología Celular, Facultad de Medicina, UCM, Pl. de Ramón y Cajal, s/n, 28040 Madrid, Spain; Charité – Universitätsmedizin, Corporate member of Freie Universität Berlin, Humboldt-Universität zu Berlin, Berlin Institute of Health, Department of Neurology with Experimental Neurology, Charitéplatz 1, Berlin 10117, Germany; Charité – Universitätsmedizin, Corporate member of Freie Universität Berlin, Humboldt-Universität zu Berlin, Berlin Institute of Health, Institute for Medical Immunology, Augustenburger Platz 1, Berlin 13353, Germany; Charité – Universitätsmedizin, Corporate member of Freie Universität Berlin, Humboldt-Universität zu Berlin, Berlin Institute of Health, Neuroscience Clinical Research Center, Charitéplatz 1, Berlin 10117, Germany; Charité – Universitätsmedizin, Corporate member of Freie Universität Berlin, Humboldt-Universität zu Berlin, Berlin Institute of Health, Center for Stroke Research Berlin, Charitéplatz 1, Berlin 10117, Germany; Division of Neuroscience, School of Biological Sciences, Faculty of Biology, Medicine and Health, The University of Manchester, Manchester, UK; Geoffrey Jefferson Brain Research Centre, Manchester Academic Health Science Centre, Northern Care Alliance NHS Foundation Trust, University of Manchester, Manchester, UK; Departments of Neurology, Psychology and Neurosurgery, Arizona Center on Aging, and BIO5 Institute, University of Arizona, Tucson, AZ, USA; Department of Neurosurgery, Stanford University School of Medicine, Stanford University, Stanford, CA, USA

**Keywords:** Cognitive behavioral testing, multi-center trial, neurofilament light chain, post-stroke cognitive impairment, stroke

## Abstract

Cognitive decline is a significant long-term consequence of stroke and has no available treatments. To aid in therapy development, we sought to achieve robust detection of cognitive performance after stroke in a multi-site design. Ischemic stroke was induced in adult and middle-aged male C57BL/6J mice utilizing three well-established models: distal middle cerebral artery occlusion (dMCAO), dMCAO with hypoxia and transient MCAO. Cognitive outcomes were assessed via Novel Object Recognition (NOR) and Barnes Maze (BM) tests prior to surgery, and during sub-acute (1-2 weeks) and chronic (8 weeks) phases post-stroke. Histology and immunostaining were used to assess infarct size, tissue damage and neuronal loss, and plasma neurofilament light was quantified. We did not detect a reliable cognitive deficit after stroke using NOR but saw a promising signal from BM (single site tested only). Overall, our study highlights the often-encountered challenges in detecting post-stroke cognitive impairment within the pre-clinical stroke community, as well as a number of complexities in the design and execution of pre-clinical stroke cognition studies, particularly as applied to a multi-site structure. We provide recommendations and suggest important aspects of stroke cognition studies to consider in the future, whether operating as an individual lab or a multi-site group.

## Introduction

Stroke represents a significant global health burden, contributing to high rates of mortality and disability. Post-stroke cognitive impairment (PSCI) has been identified by survivors as one of the most distressing consequences. PSCI affects up to one-third of stroke survivors within five years, and greatly diminishes their quality of life.^1–3^ This is underscored in surveys conducted among both patients and healthcare professionals, which identified the foremost priority in the field to be research aimed at improving cognitive symptoms such as impairments in memory and concentration.^4–6^ Notably, stroke doubles the risk of subsequent cognitive impairment, irrespective of controlling for established vascular risk factors such as hypertension and obesity, or the prevention of additional infarcts.^2^ PSCI represents a critical unmet need due to the absence of available treatments and lack of mechanistic understanding.^2,6,7^

One way to address this unmet need is through the development and characterization of pre-clinical stroke models that generate robust cognitive deficits. Such models would create opportunities for pre-clinical screening of candidate interventions and improve prospects for translation into clinical therapies.^8,9^ Previous networks have conducted therapeutic testing in multi-center, randomized pre-clinical trials utilizing stroke size as a primary endpoint.^10–12^ The Stroke Preclinical Assessment Network (SPAN) has made important advances by establishing a multi-center trial protocol with sensorimotor function changes as the primary readout.^13,14^ Nonetheless, a need still remains to consider other functional outcomes relevant to stroke survivors, including cognitive decline, altered mood and fatigue. Researchers could then validate potential therapeutics using robust outcome measures with high clinical relevance.^15–18^

Rodents have a remarkable ability for spontaneous recovery after ischemic stroke, making long-term behavioral testing challenging.^15^ Cognitive testing, in particular, is time intensive, can be hard to interpret and is difficult to perform reproducibly. Additionally, inter-lab inter-investigator and inter-animal variability are all difficult to overcome and present a major challenge for pre-clinical multicenter trials. Due to these challenges which are well-recognized within the field, there is likely an inherent publication bias against negative or neutral results from these studies, as observed generally in life science research, limiting the ability of the field to identify best practices.^19,20^

It is also important to consider that cognitive decline after stroke likely arises from multiple etiologies. Acute cognitive decline is directly related to lesion size and location.^6,21^ While these direct effects may persist, additional mechanisms such as neurodegeneration and/or unresolved harmful inflammation may also underlie new diagnoses of PSCI in the chronic phase.^22,23^ Altered brain-wide connectivity is also likely a factor in longer-term cognitive changes after focal stroke.^23^ Thus, it is important that cognitive tests can reliably detect a deficit at multiple timepoints after stroke, including chronic ones. Additionally, human ischemic stroke is a heterogeneous and complex disease, due to high inter-patient variation, which cannot be fully recapitulated using any one model of pre-clinical stroke.^9,24^ Thus, assessing cognition in multiple stroke models that generate different amounts and anatomical patterns of damage will increase the sensitivity for detection of PSCI.

The Stroke-IMPaCT Network, a multi-site network comprising six labs across Europe and North America, was established to understand immune mechanisms contributing to PSCI, and from this, to identify and test novel immunomodulatory interventions. An enabling primary aim was to identify protocols that robustly detect cognitive impairment after experimental stroke in multiple labs. Our multi-site project design tested cognitive behavioral outcomes during both sub-acute (1-2 weeks) and chronic (8 week) phases of ischemic stroke. We included three stroke models, and tested at two different ages with two cognitive assays to more accurately reflect the complexity of the human stroke population. While we did not identify a combination of stroke model and cognitive test for robust detection of PSCI, we expect that our findings will aid the field (1) in managing expectations for detection of pre-clinical PSCI using commonly-applied approaches, and (2) as a basis for refining methodology to reproducibly detect cognitive impairment in pre-clinical models of ischemic stroke in future trials.

## Materials and Methods

Detailed material and methods are available in the Supplemental Materials.

### Animals

All animal protocols were performed and reported in conformity with the Animal Research: Reporting of In Vivo Experiments (ARRIVE) guidelines. Experimental procedures were approved by the appropriate animal care and use committees at each institution, and were performed under relevant national and institutional rules, including personal and project licenses. A total of *n*=150 adult (12-15 week old) and *n*=75 middle-aged (10-12 month old) male C57BL/6J mice were used (Figure 1). Sham and naïve littermates were used as controls, and all mice were randomized among experimental cages. Exclusion criteria for removing animals from behavioral testing analysis due to mortality, inability to perform cognitive tasks or tissue pathology are outlined in Figure 1B. Additional details are available in Supplemental Materials.

**Figure 1.**
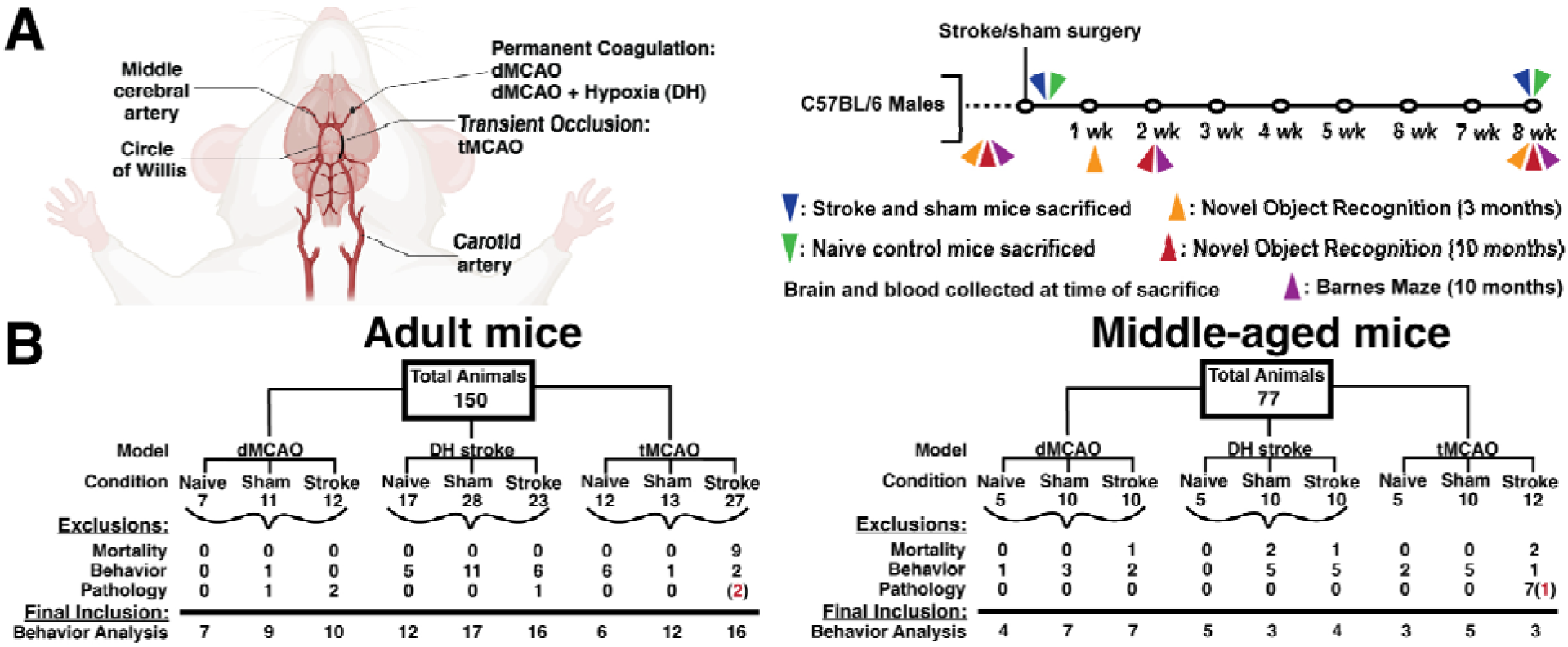
Study design. **(A)** From left to right, visual representation of points of permanent (dMCAO, DH) or transient (tMCAO) middle cerebral artery occlusion, representing the three pre-clinical models of stroke used. Study design with respective behavioral testing timepoints and times of sacrifice. This panel was created with BioRender. **(B)** Visual representation of the total number of animals used in two behavioral testing cohorts. Numbers represent the animals used in each surgical (stroke or sham) or naïve group in adult (12-15 wk) and middle-aged (10-12 mo) cohorts of male C57BL/6J mice. Animals were excluded due to surgical mortality, lack of novel object preference at baseline (Discrimination Index<0.1), or irregular pathology. Inconsistent pathology is defined as lack of stroke pathology in MCAO animals, or aberrant pathology present in sham or naïve animals. Red numbers represent the number of animals excluded from all analyses for both pathology and task performance. dMCAO: Distal middle cerebral artery occlusion; DH: dMCAO + Hypoxia stroke; tMCAO: transient middle cerebral artery occlusion.

### Stroke Models

Three pre-clinical stroke models were used in this study: proximal transient (filament) middle cerebral artery occlusion (tMCAO) was performed at Weill Cornell Medical College (WC; 25 min occlusion) and Charité Universitätsmedizin Berlin (CUB; 60 min occlusion), permanent distal middle cerebral artery occlusion (dMCAO) was performed at Stanford University (SU), University of Edinburgh (UoE) and Centro Nacional de Investigaciones Cardiovasculares (CNIC), and dMCAO + Hypoxia (DH) surgery was performed at University of Arizona (UoA; 40 min hypoxia) and SU (60 min hypoxia), as described previously.^22,25–31^ Site specific surgical procedures are outlined in Supplemental Materials.

### Behavioral Testing

#### Novel Object Recognition

Novel Object Recognition (NOR) was performed on adult and middle-aged cohorts, as previously described.^32–34^ Site-specific differences in NOR protocols used for the adult (12-15 week old) cohort are available in Table 1, while a single unified NOR protocol was used by all sites for the middle-aged (10-12 month old) cohort. Briefly, the unified NOR task consisted of two phases: a training phase (Phase I) and testing phase (Phase II). In Phase I, a single animal was placed in the arena with two identical objects and allowed to explore freely for 5 min. Phase II started 3 h after the completion of Phase I. Here, mice were exposed to one familiar and one novel object for 5 min, and their interactions with the objects were recorded. Objects were counterbalanced between animals, to avoid any innate object preference. Time spent interacting with each object was manually recorded by a blinded experimenter. To assess task performance, a discrimination index (DI) score was calculated with: (t_novel_-t_familiar_)/(t_novel_+t_familiar_). A score closer to −1 indicates a preference for the familiar object, a score closer to +1 indicates a preference for the novel object, and a score close to 0 demonstrates no object preference.

**Table 1.** Laboratory specific differences in NOR protocols. Adult (12-14 wks old) male C57BL/6J mice were tested on the Novel Object Recognition (NOR) protocol prior to, and 1 and 8 weeks after sham or stroke surgery. Each site had minor variations in NOR testing protocol.

| Site | Handling Time | Habituation Time | Room Lighting | Material & Size of arena | Inclusion & exclusion cutoff used? | Other notes |
| --- | --- | --- | --- | --- | --- | --- |
| UoA | 3 days prior to testing, 5 min per mouse per day | Mice in the room 1 hr before testing | Dim | Light gray acrylic<br>40cm (W),<br>40cm (L),<br>35cm (H) | Must interact with both objects during phase 1<br>Drilling noise during testing led to removal of 4 sham animals at baseline timepoint | No white noise<br>Temperature controlled room.<br>Location of novel object was switched for half of the mice to prevent location bias |
| SU | 3 days prior to testing, 20 min per mouse per day | Mice in the room 1 hr before testing<br>To arena: 5 min prior to phase 1 | Dim/dark | Blue acrylic<br>20in (W),<br>20in (L),<br>18in (H) | No, all animals were included in the study | White noise playing in background during testing |
| UoE | 3 days prior to testing, 5 min per mouse per day<br>(Note: for 8-week timepoint mice were handled again) | 2 days habituation prior to testing:<br>Day 1: 30 min exploration with cage mates, 30 min for each individual mouse<br>Day 2: 20 min exploration with cage mates, 20 min for each individual mouse | Dim | Dark grey acrylic<br>49.5cm (W),<br>50cm (L),<br>25cm (H) | No, all animals were included in the study | No white noise<br>Temperature controlled room |
| WC | None beyond testing | Mice in the room 1hr before testing<br>To arena: 5 min exploration 1 day before test day | Dark room, bright lights around arena | Transparent acrylic surrounded by white construction paper on the outside<br>47cm (W), 30cm (L), 29cm (H) | For baseline, mice had to display preference for novel object. No criteria applied after baseline | Room is subject to temperature swings and vibration |
| CUB | 3 days prior to testing, 20 min per mouse per day | Mice in the room 1hr before testing<br>To arena: 5 min prior to phase 1 | Dim | Transparent acrylic<br>45cm (W), 45cm (L), 14.3cm (H) | No, all animals were included in the study | No white noise<br>Temperature controlled room |

#### Barnes Maze

A modified Barnes maze (BM) protocol was utilized as previously described,^32,33^ with minor modifications for middle-aged mice. Briefly, the BM consisted of a large, circular platform with 16 holes on the outer edge, which was positioned approximately 90 cm off the floor in the center of the room. All maze holes were left open to the floor, except for one which contained an escape hole. The escape hole was aligned with one of four visual cues, which were equally spaced around the testing room. Mice performed four trials per day for four consecutive days at each testing timepoint. The escape hole position was fixed for the entirety of the task, while the starting position of each trial within a day was altered relative to the escape hole position. Each trial had a maximum duration of 90 s. If a mouse failed to find the escape hole within 90 s, it was gently guided towards the escape hole by tapping its tail and hind legs. The time (in seconds) to identify the escape hole (primary latency), defined as a mouse entering the hole opening with at least their head, was manually recorded.

### Tissue Collection and Processing

Mice were euthanized at 3 or 56 days following stroke by exsanguination and intracardiac perfusion with 0.9% saline (without anticoagulant) under isoflurane anesthesia (UoE, UoA), sevoflurane (CNIC), sodium pentobarbital (WC), or xylazine/ketamine (SU, CUB). Brain tissues and blood were collected. Brains were extracted and post-fixed in 4% paraformaldehyde (PFA) in phosphate buffer for 24 h, then transferred to 30% sucrose solution (with 0.1% sodium azide) until sectioning. All brain tissue was sectioned at 40 μm thickness with a freezing microtome (UoA, SU) or cryostat (UoE), and sequentially collected into 16 tubes with cryoprotective medium (30% glycerin, 30% ethylene glycol, 40% 0.5 M sodium phosphate buffer). All sections were stored at −20°C in cryoprotective media.

Blood was drawn from the vena cava or left ventricle using an EDTA-impregnated syringe. Microvette 500 K3EDTA tubes (Sarstedt #201341) containing whole blood were centrifuged at 4°C for 10-15 min at 1,000 g. Subsequently, a second centrifugation for 15 min at 2,000 g was performed to deplete platelets. The resulting supernatant was designated as plasma and stored at −80°C until use.

### Histopathology and Immunostaining

#### Hematoxylin & Eosin (H&E) Staining

All H&E and immunofluorescence staining was carried out at UoE. H&E reagents were purchased from Epredia (UK, Instant Hematoxylin #12687926) and Cell Path (UK, Eosin Y #RBC-0201-00A). Sections were washed in PBS at room temperature (RT) to remove cryoprotectant. Sections were then mounted, washed in dH_2_O, and dried overnight prior to H&E staining, using a standard protocol.^35^ Briefly, sections were dipped 10 times in 95% EtOH and rinsed in dH_2_O for 30 s. Slides were incubated in hematoxylin for 1 min 30 s and then rinsed in dH_2_O until the water ran clear. Slides were then dipped in acid alcohol for 30 s and rinsed in dH_2_O for 30 s. Next, slides were dipped into Scott’s tap water for 1 min and rinsed in dH_2_O. Then, slides were dipped once in eosin and washed in dH_2_O. Finally, slides were dehydrated in increasing concentrations of EtOH, cleared in xylene and mounted in DPX mountant.

H&E staining was performed on two sections per animal: one rostral (Bregma −0.70 mm to −0.82 mm) and one caudal (Bregma −1.70 mm to −2.18 mm). Percent infarct area was calculated at 3 days and 56 days on the rostral brain section only using the Swanson method,^36,37^ as this brain level best captures the stroke lesion across all three models. The regional distribution of gross pathology was quantified on both rostral and caudal sections (ROIs: rostral cortex, caudal cortex, thalamus, hippocampus). Pathological evidence of stroke was defined on H&E as presence of vacuolation, and/or pallor, and/or neuronal perikaryal shrinkage within each ROI (Figure 2C, Supplementary Figure 4). We also quantified ventricular enlargement (a proxy for brain atrophy) and displacement of the corpus callosum. If pathology was identified within a defined ROI, the animal was given a score of 1 for that metric, and if no damage was identified the score was 0. We then calculated sums to identify the number of animals per model that exhibited damage in each region.

**Figure 2.**
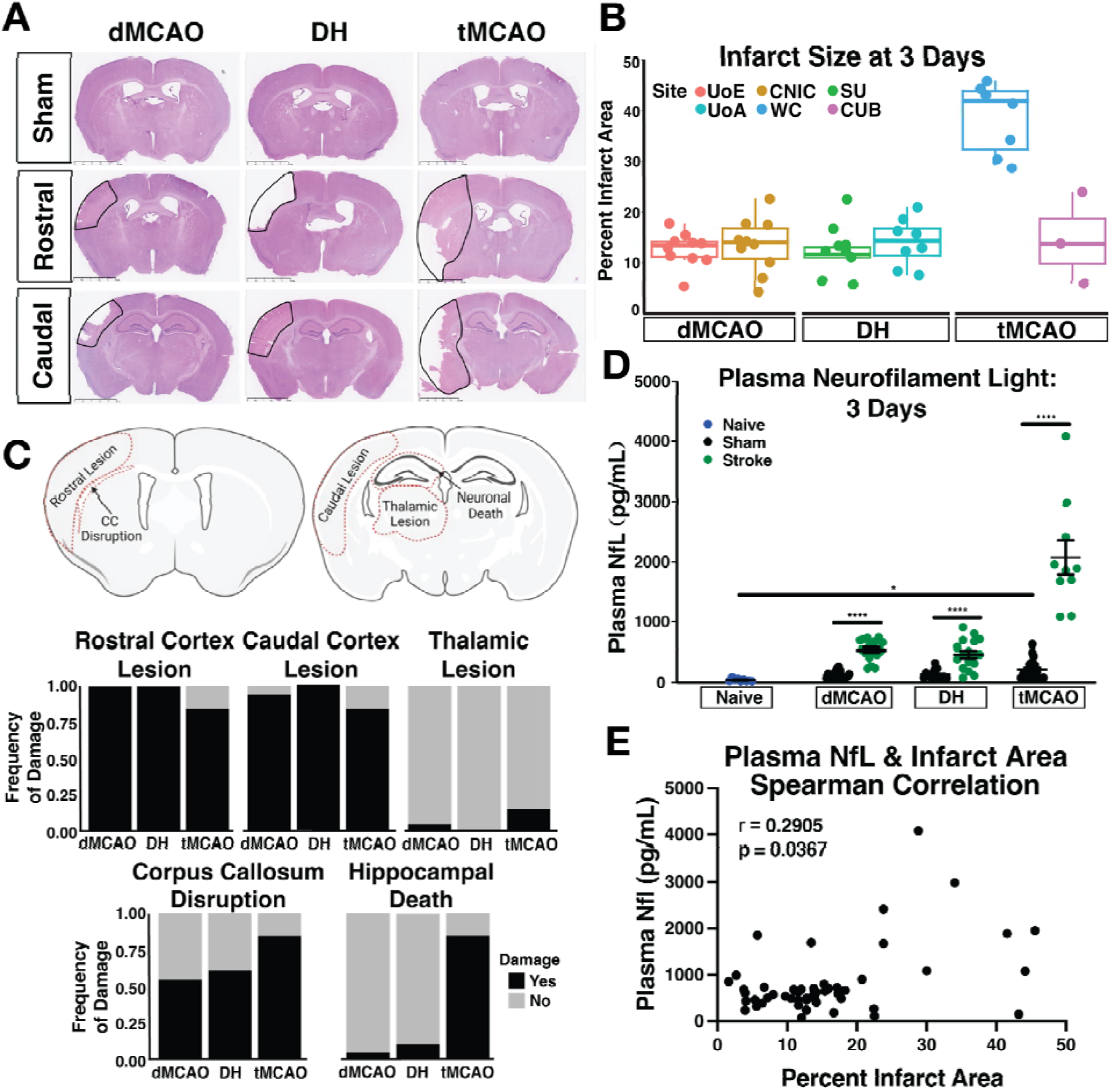
Acute (3 days) histopathology assessment in 3 pre-clinical stroke models. **(A)** H&E representative images from each stroke model (dMCAO, DH, tMCAO) at rostral (Bregma −0.70mm to −0.82mm) and caudal (Bregma −1.70mm to −2.18mm) brain levels. The infarct is outlined with a solid black line. Scale bar = 2.5mm **(B)** Quantification of infarct size at 3 days after dMCAO, DH or tMCAO across the six sites. **(C)** Anatomical locations for assessments of pathological damage frequency. Assessment of damage was conducted on H&E staining and scored categorically as absent (score of 0) or present (score of 1) in each anatomical location. A frequency score of 1 indicates that all animals within that group and at that anatomical location showed pathological damage. **(D)** Plasma NfL concentration in each stroke model 3 days post ischemia, and respective sham controls. Naïve mice were analyzed from SU only. **(E)** Plasma NfL concentration correlation with percent infarct area in a Spearman correlation analysis. (Naïve: *n*=9; dMCAO: *n*=20 per grp; DH: *n*=17-18 per grp; tMCAO: *n*= 10-17 per grp). Data are presented as mean ± SEM and statistics represent a two-way (B, D) ANOVA with Tukey’s post hoc test \**p*<0.05; \*\*\*\**p*=<0.0001. dMCAO: Distal middle cerebral artery occlusion; DH: dMCAO + Hypoxia stroke; tMCAO: transient middle cerebral artery occlusion.

#### NeuN Immunofluorescence

Immunofluorescent staining followed a standard protocol. Briefly, sections at Bregma level −1.34 to −2.06 were washed in PBS at RT. Sections were then blocked for 2 h (10% Donkey serum in PBS-T), before overnight primary antibody incubation with NeuN (Abcam, ab177487, 1:1000; 10% Donkey serum in PBS-T) at 4°C. Next, sections were incubated with a fluorophore-conjugated secondary antibody for 2 h at RT. Sections were then washed, counterstained with DAPI and mounted in Prolong Glass Anti-Fade (Thermofisher, #P36984). Percent coverage of NeuN was calculated by thresholding hippocampal ROIs in QuPath (v0.5.1).

### Evaluation of Plasma Neurofilament Light (NfL)

NfL was quantified using the Simoa® NF-Light v2 Advantage Assay (Quanterix®, Cat. No. 104073) according to manufacturer instructions. Plasma samples, controls, and calibrators were measured in duplicate. The two-step immunoassay uses paramagnetic beads coated with anti-NfL antibodies to capture target molecules, which are then detected by biotinylated antibodies and labeled with streptavidin-ß-galactosidase. Signal detection was performed using the Simoa® optical system after adding substrate solution and transferring the beads to the Simoa® disc.

### Statistical Analysis

Experimenters were blind to surgery status throughout data collection and analysis. Statistical testing was performed in R (version 4.4.2) or Graphpad Prism 10. Normality was assessed with a Shapiro-Wilk test. Histological and immunofluorescence data were analyzed with one-way ANOVA with Tukey’s post-hoc test. Baseline behavioral data were analyzed with one-way ANOVA with Tukey’s post-hoc test, and behavioral performance over time was analyzed with a two-way repeated measures ANOVA with Tukey’s post-hoc test. Statistical variance was assessed with f-test for equal variance, manual and automated NOR scoring was compared with the Bland-Altman method comparison, infarct size was correlated to plasma NfL concentration with a Spearman correlation analysis, and inter-rater variability was assessed with the intraclass correlation coefficient. For all statistical testing, *p*<0.05 was considered statistically significant.

## Results

### Study Design and Exclusions

We implemented our study across five sites using three different stroke models: permanent distal middle cerebral artery occlusion (dMCAO), dMCAO with hypoxia (DH), and transient filament MCAO (tMCAO; Figure 1A). Size and neuroanatomical distribution of the primary damage (infarct and associated neuronal death) may impact cognition in both the acute and chronic phases. Indeed, individual network laboratories have previously detected cognitive deficits when testing these same models at similar timepoints.^22,23^ With differing size/distributions produced by the different models (and potentially similar models applied across different sites) anticipated, this would generate useful heterogeneity, and reveal whether certain models and associated pathological patterns may be more strongly associated with PSCI. To assess cognitive impairment after stroke in adult mice, we utilized the NOR test. NOR is one of the most commonly used cognitive tests in rodents, has low time commitments and was readily implemented at all sites at low cost.^34^ We tested cognitive outcomes in both sub-acute (1-2 weeks) and chronic (8 week) phases of ischemic stroke in adult (12-15 week old) and middle-aged (10-12 month old) mice. The BM test was introduced in the middle-aged cohort to assess spatial working and reference learning/memory (Figure 1A). For adult behavioral analyses, 45 mice were excluded from analysis due to surgical mortality, no preference for the novel object at baseline (discrimination index, DI<0.1) or lack of stroke pathology on histopathological examination, as outlined in Figure 1B. Similarly, 33 animals were excluded from the middle-aged cohort. Surgical mortality primarily affected mice assigned to the tMCAO stroke group, and a lack of novel object preference at baseline was a contributor to lower sample size across all sites and ages.

### Stroke Model Comparative Pathology

We first assessed the patterns of acute stroke pathology to investigate model consistency across sites, and to establish patterns of pathological damage. A cohort of *n*=40 dMCAO, *n*=44 DH and *n*=32 tMCAO adult (12-15 week old) male C57BL/6J mice compiled from all six sites was euthanized 3 days after surgery. Edema-corrected infarct measurements 3 days after stroke revealed that the dMCAO and DH models result in consistent infarct sizes (∼12% of hemisphere) between sites (Figure 2A, B). In contrast, infarct size at sites using the tMCAO model was more variable. Infarct size at WC was 45% of the hemisphere, while at CUB it was 15% (Figure 2B). Infarct size in the tMCAO model was unrelated to the duration of occlusion, likely reflecting the model’s inherent variability. Infarcts generated by this model are impacted by blood flow through the Circle of Willis, which is anatomically variable in C57BL6 mice.^38,39^

We next determined the neuroanatomical distribution of primary stroke pathology by assessing the presence or absence of pathological damage at five distinct anatomical locations in rostral and caudal sections (Figure 2C). All three models had significant cortical damage in both rostral and caudal sections, with minimal thalamic damage. In contrast, the tMCAO model had an elevated frequency of hippocampal death compared to either the dMCAO or DH models.

We next measured plasma NfL, an emerging biomarker for neurodegeneration in aging and Alzheimer’s disease, to assess the relationship with primary stroke damage.^40–42^ In all three models, plasma NfL concentration was significantly elevated 3 days after stroke compared to sham animals. There were no significant differences in plasma NfL concentration in naïve mice compared to dMCAO or DH sham animals, while the tMCAO-sham animals had moderately elevated plasma NfL compared to naïve animals. Consistent with a larger infarct size and a higher frequency of damage across anatomical areas in the tMCAO stroke samples, the concentration of plasma NfL was also significantly elevated compared to the dMCAO and DH models (Figure 2D). Spearman correlation analysis between plasma NfL concentration and stroke size (Figure 2E) show that plasma NfL correlates moderately with infarct size at 3 days after stroke (Spearman rho=0.2905; *p*=0.0367).

We next assessed chronic pathological damage at 56 days post-stroke given the importance of chronic neurodegeneration to cognitive impairment. Infarct size in all models was largely similar to the 3 day timepoint (Figure 4A, B), although it should be noted that the histopathological appearance of the infarcts was substantially different (e.g. signs of dense inflammatory cell infiltrates), as expected. Marked hemispheric atrophy of the ipsilateral hemisphere was common, as reflected by a high frequency of ventricular enlargement (Figure 4C). The distribution of overt pathology at 56 days (Figure 4C) was broadly similar to 3 days. There was a minor reduction in the frequency of caudal cortical damage in the dMCAO model, likely reflecting infarct resorption, and more frequent thalamic pathology in the tMCAO model, potentially due to secondary remote degeneration. We also assessed NeuN coverage as a marker for neuronal death in the hippocampus, however saw no reductions in NeuN coverage in any model after stroke (Figure 4D, E). In contrast to the acute timepoint, at 56 days there was no difference in plasma NfL concentration in dMCAO or DH stroke mice compared to their respective shams (Figure 4F). We did, however, observe elevated plasma NfL in tMCAO stroke mice compared to shams. Overall, concentrations of plasma NfL were markedly lower at 56 days after stroke compared to the 3 day timepoint in all groups, and plasma NfL was no longer correlated with stroke size at this chronic timepoint (Spearman rho=0.1289; *p*=0.3220). Overall, these data show the early and chronic pathological characteristics associated with each of the models revealing both inter-site/model consistencies and useful inter-model heterogeneity in infarct size/distribution, as well as acute-to-chronic evolution for evaluation of cognitive changes.

### Subacute and Chronic Cognitive Assessments in Adult Mice

We first determined whether inter-rater variability in the NOR task was sufficiently low to compare assessments from different sites, by randomly selecting two videos from each site (1 sham and 1 stroke) for manual scoring by one assessor from each site. This resulted in five assessments per video. We determined that the intraclass correlation coefficient was 0.6991 (95% CI=0.4548-0.894; Supplementary Figure 1). Additionally, to confirm that manual assessment of the NOR task recordings was accurate compared to automated analysis with AnyMaze software, one site (UoA) utilized both analysis methods on the same set of videos. Both manual and automatic scoring methods resulted in consistent outcomes, and the differences between methods were not dependent on the final DI value (i.e. neither method was better at detecting preference for a given object; Supplementary Figure 2). Given these results and the cost of purchasing AnyMaze software across all sites (which is also relevant when considering utility to deploy across the research community), each site manually analyzed their own videos while blinded to experimental status.

We first analyzed baseline performance at individual sites to establish whether all sites had a working task at baseline (Supplementary Figure 3). This revealed that half of the sites had a high percentage of mice with no novel object preference (DI<0.1). We therefore evaluated site-specific environment and protocol differences. While certain differences that impact performance, such as anxiety and noise, are unable to be quantified, we documented quantifiable differences such as handling and habituation time in Table 1. We hypothesized that handling and habituation time may impact NOR performance, independent of any surgical intervention. We therefore categorized sites based on none (WC), low (UoE, UoA) or high (SU, CUB) handling, and none (UoA), low (SU, CUB) or high (UoE, WC) habituation times (Figure 3B, C). The mean DI was unaffected by the amount of handling (Figure 3B). In contrast, the mean DI was significantly higher at sites with high habituation compared to those with low or no habituation (Figure 3C). We next analyzed the variance in DI at different sites to determine whether differences in handling or habituation may impact overall variability and subsequent statistical power. While certain sites had significantly lower variability in DI (SU, WC) compared to others (UoA, UoE, CUB), there was no statistical relationship between the amount of handling and habituation time and the variability in DI measures (Supplementary Figure 3). We therefore established a uniform baseline for exclusion across sites: a baseline DI score of <0.1. This led to the exclusion of 14% of tested mice (Figure 3D). We chose a DI=0.1 which is permissive to inclusion while still maintaining a degree (>10% more time) of novel object preference (>0).

**Figure 3.**
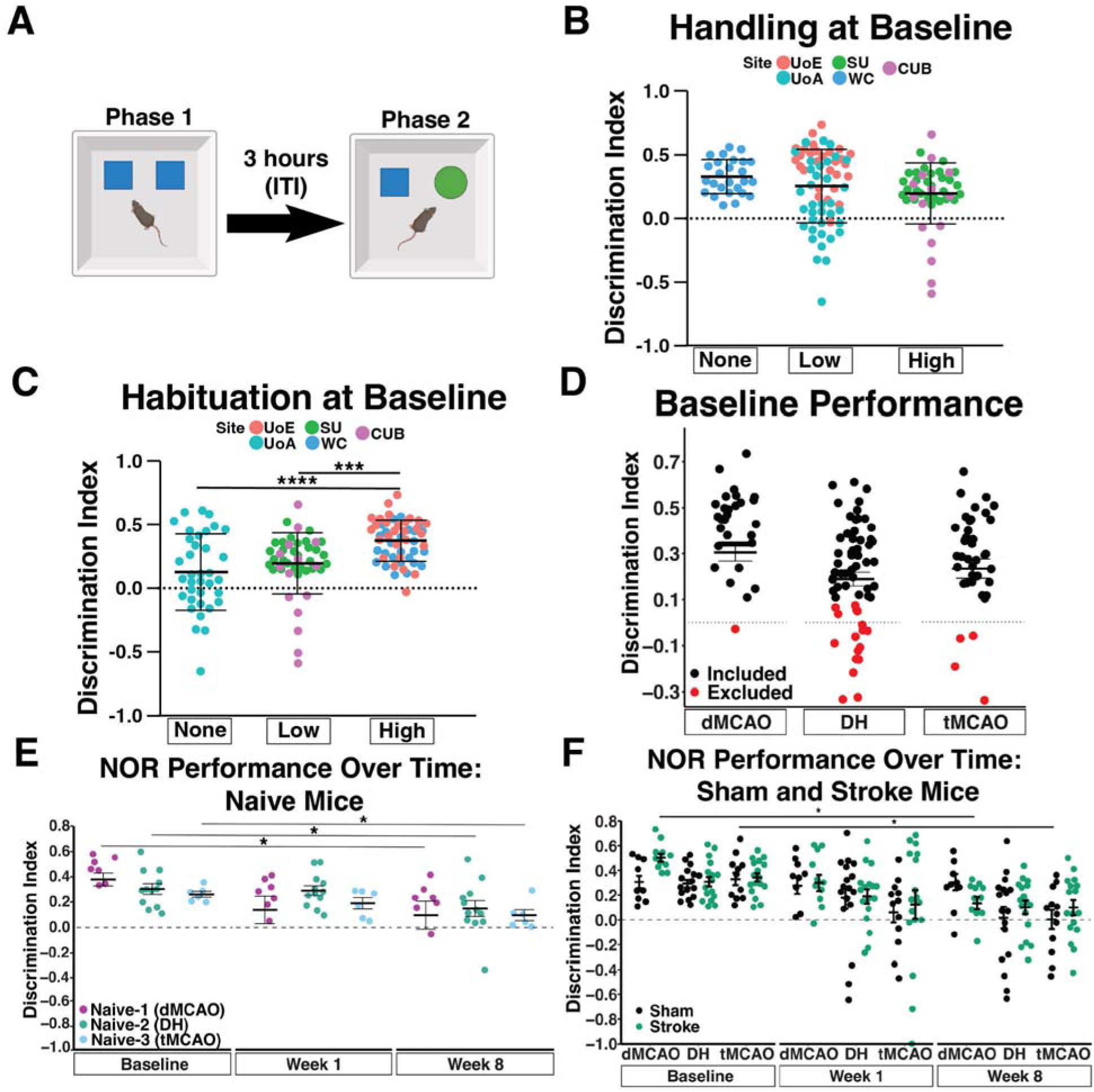
Adult mouse performance on Novel Object Recognition (NOR) task. **(A)** Schematic of the NOR test; 12-15 wk old male C57BL/6J mice are presented with two identical objects in Phase 1, and after 3 h (inter-trial interval; ITI) one familiar object was swapped with a novel object (Phase 2). Interaction time for each object was recorded to calculate a Discrimination Index (DI). **(B)** Average DI at individual sites, categorized by the amount of handling **(B)** or habituation **(C)** (See Table 1 for details) performed at baseline. (WC, *n*=24; UoE, *n*=26; UoA, *n*=23; SU, *n*=29; CUB *n=*6). **(D)** DI of all mice assigned to each stroke model (dMCAO, DH or tMCAO) prior to surgical intervention (baseline performance). A DI<0.1 merited exclusion for lack of novel object preference. Animals excluded from behavioral analyses are plotted in red. The dotted line represents DI=0, or no object preference. **(E)** DI of all naïve mice assigned to a surgical model at baseline, week 1 and 8 as a measure of performance after repeated testing. (dMCAO, *n*=7; DH, *n*=12; tMCAO, *n*=6). **(F)** DI score in the NOR task of sham and stroke mice for each surgical model. (dMCAO, *n*=9-10 per grp; DH, *n*=16-17 per grp; tMCAO, *n*=12-16 per grp). All data are presented as mean ± SEM, and were analyzed by one (C-E) or two-way (F) repeated measures ANOVA with Tukey’s post hoc test \**p*<0.05; \*\*\**p*<0.001; \*\*\*\**p*<0.0001; dMCAO: Distal middle cerebral artery occlusion; DH: dMCAO + Hypoxia stroke; tMCAO: transient middle cerebral artery occlusion.

**Figure 4.**
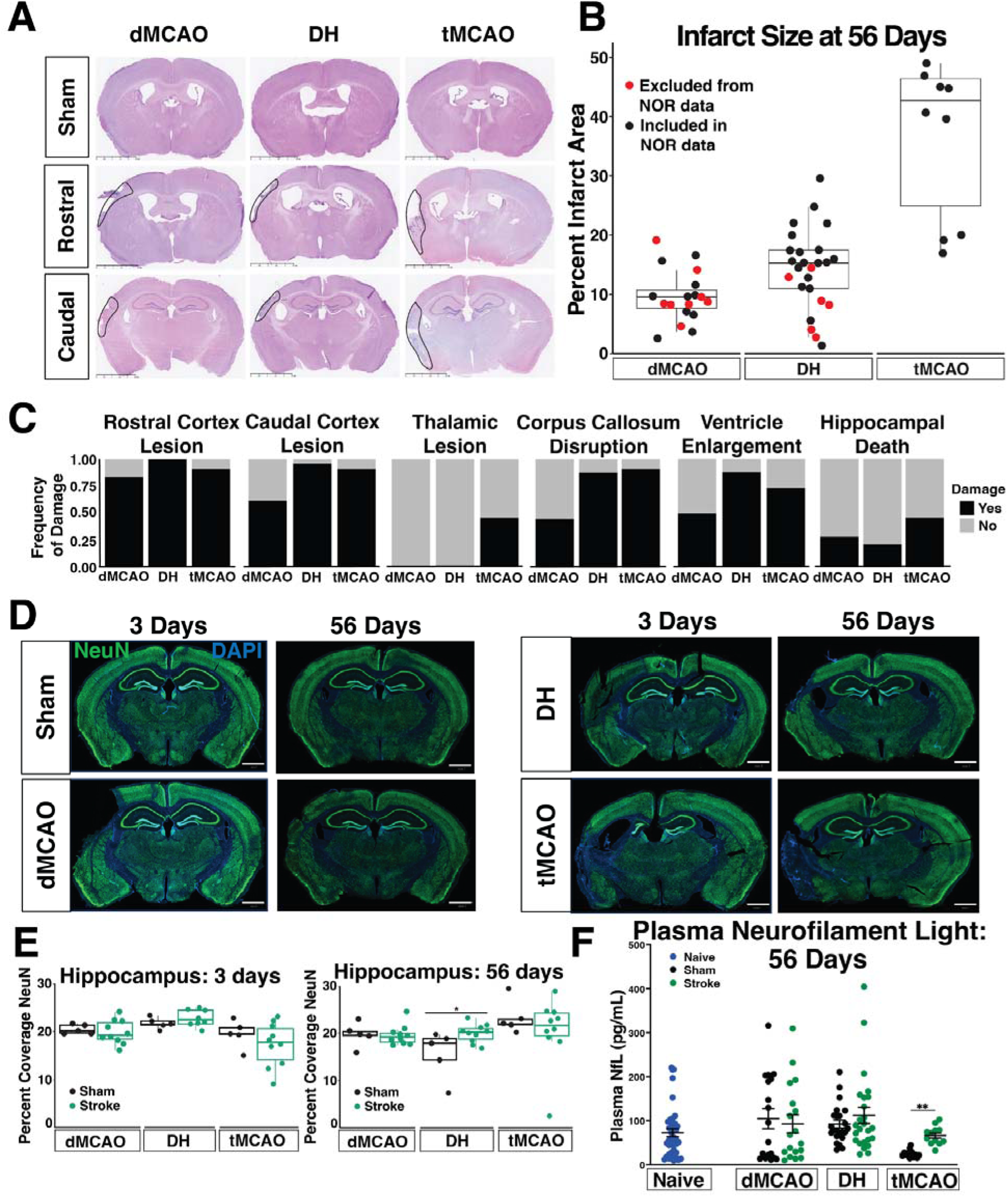
Histopathology assessment at 56 days after stroke in adult mice. **(A)** H&E representative images from each stroke model (dMCAO, DH, tMCAO) at rostral (Bregma −0.70mm to −0.82mm) and caudal (Bregma −1.70mm to −2.18mm) brain levels. Scale bar=2.5mm. Infarcted tissue is outlined with a solid black line. **(B)** Quantification of infarct size, at 56 d post-surgery. Red circles represent mice that have been used for histopathological assessment but were not included in NOR analysis (dMCAO, *n*=18; DH, *n*=25; tMCAO, *n*=9). **(C)** Frequency of damage distribution in dMCAO, DH and tMCAO stroke models on H&E stained sections. **(D)** Representative images of whole brain NeuN staining at 3 or 56 days after sham or stroke surgery. Scale bar=1mm **(E)** Quantification of NeuN percentage coverage in the hippocampus. (dMCAO, *n*=5-10 mice/group; DH, *n*=5-8 mice/group; tMCAO, *n*=5-10 mice/group). **(F)** Plasma NfL quantification in each stroke model 56 d post ischemia and respective sham controls. Naïve mice (*n*=13-19/model) were pooled. (dMCAO, *n*=18-20 per grp; DH, *n*=25-26 per grp; tMCAO, *n*=9-15 per grp). In all panels, data are presented as mean ± SEM and analyzed by one or two-way ANOVA with Tukey’s post hoc test \**p*<0.05; \*\**p*<0.01. dMCAO: Distal middle cerebral artery occlusion; DH: dMCAO + Hypoxia stroke; tMCAO: transient middle cerebral artery occlusion.

We next asked whether re-testing the same naïve mice affected longitudinal performance on the NOR task, as a potential confounding factor (familiarity yielding a lower DI score) for identifying cognitive impairment in stroked animals. Indeed, across all sites, naïve mice had a significantly lower DI at the 8-week timepoint compared to their respective baseline performance (Figure 3E).

Having established baseline and longitudinal NOR patterns across sites in the absence of MCAO, we next determined whether any of the models cause a stroke-specific change in NOR performance at acute and chronic post-stroke phases. We observed no differences in NOR performance between stroke and sham mice from any model, at any timepoint after surgery (Figure 3F). Similarly to naïve mice, there was a trend towards reduced DI in all stroke models when comparing DI at baseline to DI at 8 weeks after surgery, alongside increased variability.

### Subacute and Chronic Cognitive Assessments in Middle-Aged Mice

Given our inability to detect cognitive impairment after stroke in young adult mice, we next altered the design to increase sensitivity. Older mice have exacerbated cognitive problems after stroke,^21^ and we therefore hypothesized that cognitive testing may be more sensitive at older ages. Therefore, we used middle-aged (10-12 month old) mice, and also added the BM as a second cognitive test. Although BM testing requires considerably more time and experimenter skill, we hypothesized that it may be more sensitive to post-stroke deficits by probing additional cognitive domains. Also, previous researcher experience suggested it may reduce exclusions due to poor baseline performance.^32,33^ In addition, we unified our NOR protocol to eliminate the site-specific differences in handling, habituation, and object location outlined in Table 1.

Here, we found that at baseline 37% of mice had a DI<0.1 on the NOR test, and thus excluded these mice from subsequent NOR analyses (Figure 5A). We next assessed performance in stroke animals, compared to their respective sham controls at baseline, 2 weeks and 8 weeks after surgery. At the 2-week timepoint, there was a trend towards worse performance in the tMCAO stroke mice compared to the respective shams (*p*=0.078), however this trend did not extend to the 8-week timepoint. There was no timepoint after stroke where a cognitive deficit was detected in dMCAO or DH mice with the NOR task (Figure 5B).

**Figure 5.**
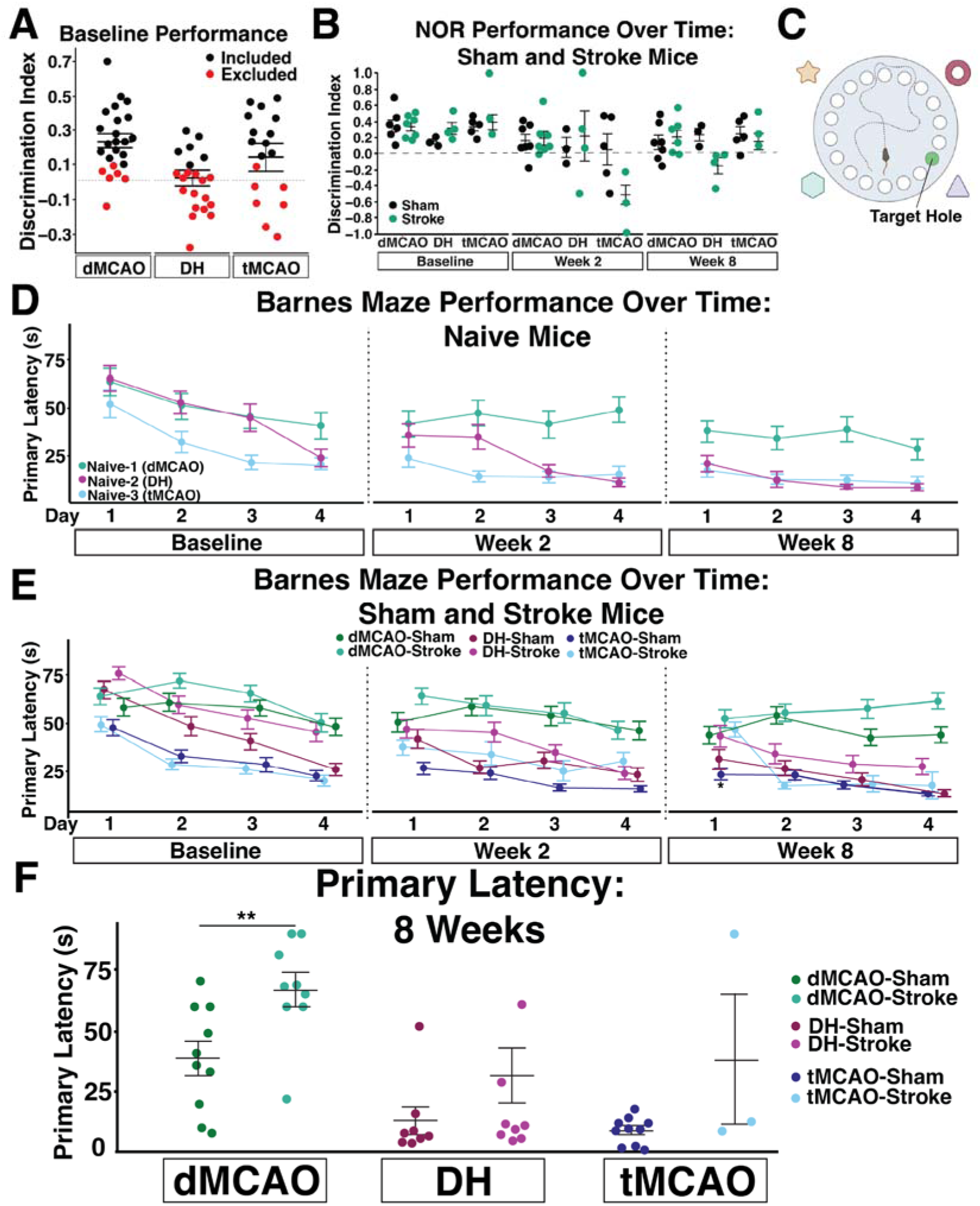
Middle-aged mouse cognitive behavior performance. **(A)** DI of all middle-aged (10-12 mo) mice, designated by assigned surgical group (dMCAO, DH or tMCAO). A Discrimination Index (DI) ≤0.1 at baseline merited exclusion from subsequent NOR analyses, which is plotted in red. **(B)** DI of sham and stroke mice separated by surgical model at baseline and weeks 2 and 8 after surgery. (dMCAO, *n*=6-7 per grp; DH, *n*=3-4 per grp; tMCAO, *n*=3-5 per grp).**(C)** Schematic of the Barnes maze (BM) test. **(D)**. BM primary latency (s) of naïve mice. (*n*=5/group) **(E)** Primary latency from Barnes maze testing of sham and stroke mice for each surgical model (dMCAO, *n*=10/group; DH, *n*=8/group; tMCAO, *n*=3-10/group). **(F)** Primary latency (s) of individual mice on the fourth trial of day 4 of testing, during the week 8 testing timepoint. BM data are presented as mean ± SEM of 4 trials per day, for 4 testing days at baseline, 2 and 8 weeks. Statistics represent results of a two-way repeated measures ANOVA with Tukey’s post-hoc test (B, D, E) or Student’s t test (F) within each surgical model. \*\**p<0.001*

In contrast to the NOR, middle-aged mice identified the escape hole in the BM at all testing timepoints with no exclusions (Figure 5D). Poor performance was defined as a mouse never identifying or entering the escape hole after four days of training. Indeed, over the course of training, the primary latency of naïve mice significantly decreases, representing improved task performance. By the 8-week timepoint, naïve mice from all sites identify the escape hole in under 50 seconds. Like the NOR task, differences in performance of naïve animals likely represent site-specific differences in the testing environment or experimenter experience.

We next assessed whether the BM could detect a cognitive deficit after stroke. At 8 weeks after stroke, the tMCAO stroke mice performed significantly worse on the first day of testing compared to their respective shams (*p*=0.0443), but performance recovered by the second day of testing (*p*=0.9249), and the mice maintained equal performance for the duration of the task. In contrast, in both the dMCAO and DH stroke models, mice performed similarly on days 1 and 2 of testing, but stroke mice trended towards worse performance on day 3 (dMCAO *p*=0.1616; DH *p*=0.3213) and 4 (dMCAO *p*=0.2313; DH *p*=0.1145; Figure 5E). Given these trends, we analyzed the performance of individual mice on the final trial of the final testing day, where we anticipated the greatest differences in performance between sham and stroke animals (Figure 5F). There was a trend towards worse performance in the DH stroke model (*p*=0.1812), and mice that underwent dMCAO performed significantly worse (*p*=0.009).

### Chronic Pathological Assessments in Middle-Aged Mice

We next assessed infarct size in each stroke model at 56 days after stroke to determine whether aging altered the extent or distribution of ischemic damage. Lesion sizes in middle-aged mice were similar to those in adult mice (Supplementary Figure 5A, B). Like adult mice, all dMCAO and DH stroke mice had rostral cortical damage, however most dMCAO mice and fewer tMCAO mice had caudal cortical damage. The presence of thalamic lesions was also specific to tMCAO mice, consistent with the adult cohort, while corpus callosum disruption occurred in fewer middle-aged dMCAO mice compared to adult mice. Additionally, there was a higher frequency of hippocampal damage in both the DH and tMCAO models in middle-aged mice, but none in dMCAO mice (Supplementary Figure 5C). Similar to adult mice, there was no difference in hippocampal NeuN coverage in any model (Supplementary Figure 5D, E).

Finally, we analyzed plasma NfL concentration to assess whether advanced age altered concentrations of plasma biomarkers of neurodegeneration. There was no significant difference in plasma NfL concentration in naïve mice from each site, and we therefore pooled them into a single group (*p*=0.112). Similarly, there was no significant difference in plasma NfL concentration between sham and stroke animals in any model at 56 days after surgery, though, like adult mice, levels were generally lower compared to 3 days.

## Discussion

Identification of pre-clinical stroke models that show robust and reproducible cognitive deficits are urgently needed to further understand mechanisms responsible for PSCI, and to test novel candidate interventions. Here, we capitalized on a multi-site infrastructure embedded within the Stroke-IMPaCT network that spans six sites in Europe and North America. Three stroke models, three testing timepoints (pre-stroke, sub-acute phase, chronic phase) and two ages were used to assess cognition after stroke, initially using NOR. We found that low baseline DI (i.e. failure to show a preference for the novel object) led to a high number of exclusions in adults, and this was exacerbated by age and less habituation time. In addition, protocol and site-specific differences contributed to high variability at some sites. Overall, NOR did not detect stroke-specific changes in cognitive performance in any model at any site at either age. We implemented the BM as a secondary measure of cognitive function, which achieved fewer exclusions and demonstrated stroke-induced cognitive changes in middle-aged dMCAO mice (single site only). Overall, our study highlights the often-encountered challenges in detecting PSCI within the pre-clinical stroke community, as well as a number of complexities in the design and execution of pre-clinical stroke cognition studies, particularly as applied to a multi-site structure. We provide recommendations and suggest important aspects of stroke cognition studies to consider in future work whether operating as an individual lab or a multi-site group.

Our multi-site structure enabled us to incorporate a greater range of stroke models than would be possible within an individual lab, while also allowing for inter-site comparisons within models. We did not aim to standardize surgical or husbandry protocols due to regulatory and operational differences across sites. This also allowed us to assess recent observations suggesting productive heterogeneity can be an asset in multi-center studies, to increase generalizability of findings. Previous multi-center pre-clinical stroke studies have had a differing purpose (i.e. to test candidate stroke therapeutics) to our current study and have adopted differing degrees of protocol harmonization across participating sites.^10–14^

We used established stroke models in the field with anticipated patterns of underlying pathology but considered it important to confirm this, to assist with interpretation of behavioral results. As expected, the distal MCAO models produced infarcts largely confined to the cortex with greater involvement of subcortical structures in the tMCAO model. The dMCAO and DH stroke models were consistent in infarct size and location across sites and surgeons, an important finding as it eliminates these factors as a source of variability. Both models are therefore good candidates for multi-center trials due to this consistency and because of low surgical time, fast recovery, and low mortality.^25^ In contrast, higher variability in the tMCAO model was observed both within and across sites. This likely reflects the inherently more variable tMCAO procedure. The stroke lesion in the tMCAO model is influenced by anatomical differences in the Circle of Willis which is highly variable in C57BL/6J mice.^25^ The lesion size in this model is also highly dependent on extent and timing of reperfusion which can vary due to the ‘no-reflow phenomenon’.^43,44^ This suggests that the tMCAO model may require additional inclusion/exclusion criteria and standardization of surgical protocols to reach a desired balance between model heterogeneity and low variability. Chronic pathology in stroke models is less well documented and more likely to be associated with cognitive function, given the potential for secondary/remote brain injury to evolve over time. The distribution of gross stroke damage remained largely consistent between acute and chronic timepoints, however more frequent pathology in thalamic and hippocampal brain regions and substantial ipsilateral brain atrophy was evident 56 days after stroke. In future studies, these structural and gross pathological changes may be captured by longitudinal MRI. Ongoing chronic neurodegeneration may also potentially be detected by blood NfL levels. We measured plasma NfL concentration in both acute (3 day) and chronic (56 day) phases. Whilst plasma NfL concentration was elevated in all stroke models at 3 days and correlated with 3 day infarct sizes, consistent with clinical data,^42,45^ by 56 days, plasma NfL was elevated only after tMCAO albeit at lower levels. Thus, while plasma NfL is a sensitive early biomarker for acute neurological damage after stroke, greater assay sensitivity may be needed to detect later degeneration, particularly in the distal MCAO models.

As in other multi-site studies, there is often a balance to strike in choosing methods with potential for widespread implementation with varying resources versus assay applicability and sensitivity. We did not detect a cognitive deficit after stroke with any variation of the NOR task. This was surprising as multiple network members have successfully used this task in the past to detect a deficit in single-site experiments.^22,46,47^ We found that with predefined object interaction criteria, NOR can be scored consistently between observers, and does not require automated testing software. Additionally, our data suggest that higher habituation increased DI score, while any amount of handling or habituation reduced the variability in DI scores. This implies that animal anxiety may play a role in outcome variability, although we can’t exclude that environmental differences at each site may also impact these observations. Finally, our results indicate that NOR is prone to re-testing effects, and thus, may not be suitable for longitudinal studies. Overall, while NOR may be a useful test to implement on an individual lab basis where protocols can be tailored to maximize performance, it is not robust for use in multi-site studies.

We observed multiple patterns of damage affecting cortical, thalamic and hippocampal areas in our three pre-clinical models, therefore it is possible that NOR is not sensitive enough to detect subtle cognitive changes. Notably, executive function and processing speed are more likely to be affected in human stroke survivors and in vascular cognitive impairment and dementia generally, suggesting that cognitive tests of memory such as NOR may not be the most sensitive for detection of PSCI. In contrast, despite low N across groups, we saw a promising signal from the BM which primarily targets spatial working memory and processing speed. Of note, we did find evidence of hippocampal neuronal perikaryal damage, suggesting that neuronal function may be impacted after stroke. This has been observed in the DH model, where hippocampal LTP is normal at one week after stroke, then progressively worsens up to 12 weeks after stroke, and is associated with a cognitive deficit.^22^ However, there was no relationship between the presence of hippocampal neuronal injury or changes in thalamic NeuN density and NOR performance in the current study. It is possible that there may be more subtle cognitive changes in the chronic phase after stroke, which require more precise cognitive testing to capture. There are many other cognitive tasks that we could not test, and we do not exclude there may be others that are effective. Where feasible, the incorporation of additional modalities such as electrophysiology, functional-MRI and calcium imaging may improve detection of changes in functional neural substrates/surrogates underpinning cognitive processes after stroke in pre-clinical models.

In addition to this, experimenter experience likely contributed to the high variability across sites for all cognitive testing, which likely impacted our ability to detect cognitive changes after stroke. Anecdotal observations across network labs have supported the idea that the ability to detect a cognitive deficit after stroke can vary by experimenter, even when other variables such as surgeon, testing environment and testing protocol are consistent. This may be due to sex differences in the experimenters, or additional intangible differences in experience and demeanor which impact the variability in outcomes of cognitive testing experiments.^48–50^ To overcome these confounding factors when testing at multiple sites, automated tasks that reduce experimenter input such as touch-screen chambers may improve detection of pre-clinical PSCI. Nonetheless, in the specific case of NOR testing here it is unlikely experimenter differences were consequential as there was a consistent lack of PSCI detected across all sites and models.

To our knowledge, we are the first network to assess cognition after stroke with a multi-site design. We were limited, however, in our ability to incorporate both sexes and additional comorbidities such as hypertension and obesity which are common in human stroke survivors.^51,52^ Mouse strain also impacts behavior and ischemic susceptibility, and incorporating additional strains may improve translatability.^53–55^ Finally, anesthetics, which have been shown to impact cognitive outcomes, varied between sites and likely contributed to increased variability within models.^58,59^ Each of these factors may increase variability in outcome measurements,^53–55^ and should be considered in future multi-site designs. Despite this, we successfully demonstrated that it is possible to reproduce consistent infarct sizes and pathological damage at multiple sites utilising the same stroke models. Another major strength of our study is the use of two cognitive tasks that assess complementary cognitive domains, and our incorporation of two age groups. Together, our results suggest that a multi-site design of this kind enables the incorporation of a greater range of stroke models and increases statistical power (via higher n), generating useful heterogeneity which would be unachievable within an individual lab.

Our findings highlight a need to manage expectations for detection of pre-clinical PSCI using commonly-applied behavioral paradigms, which should be carefully considered based on the study’s primary aims. We have provided a ranked list (Supplementary Table 2) of readily used cognitive behavior tasks and compiled our recommendations into a framework for future testing of PSCI (Figure 6). This framework was specifically designed to ensure rigor and reproducibility with a goal of improving translatability. In it, we have prioritized tasks that mice are able to perform at baseline, prior to any surgical intervention, to reduce the required animal number for the experiment and maximize statistical power. We next consider whether the task can detect a cognitive deficit after stroke at a single site, and then whether it can be replicated at an additional site with the same effect size. Finally, our framework incorporates additional refinements to reduce experimenter time and improve statistical power such as automating the task and reducing inter-rater variability. In addition, we recommend that a) all sites incorporate handling and habituation, b) repeat testing is minimized, c) exclusion criteria are defined *a priori*, d) power calculations are performed after baseline testing, prior to surgical or pharmacological intervention, and e) multiple tests are incorporated to better evaluate different cognitive domains in sub-acute and chronic phases of stroke recovery. Finally, we advise against the use of NOR in a multi-site design, particularly in aged mice, due to the high number of exclusions we observed in our study. We instead encourage future testing of the BM, as it can be successfully undertaken in older mice, is not affected by repeated testing, tests spatial and working learning/memory, and showed detection of cognitive deficits at the single site tested in our study.

**Figure 6.**
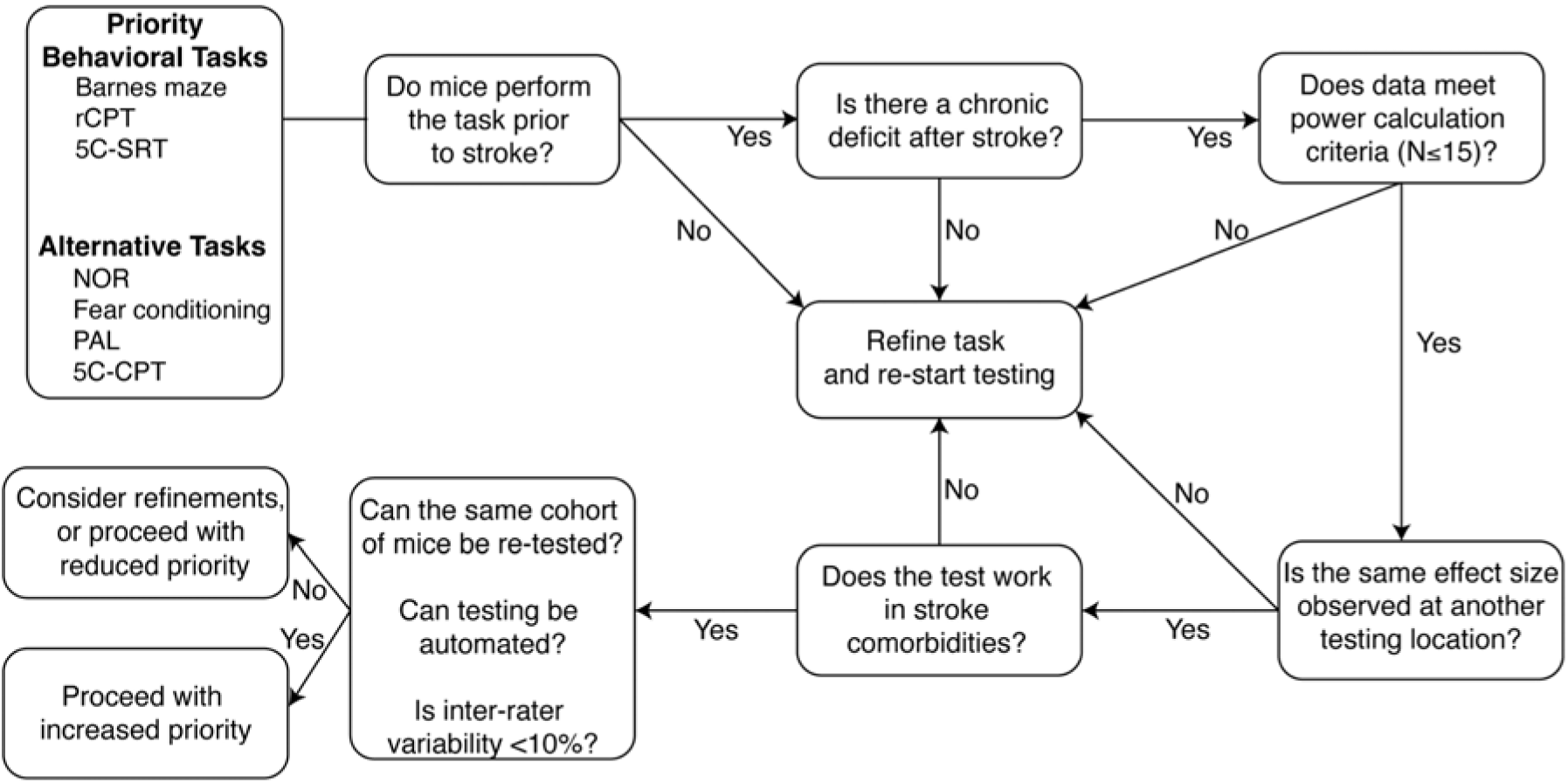
Framework for iterative testing of candidate cognitive behavioral tasks in future testing of post-stroke cognitive impairment. **I.** To ensure robust, rigorous detection of cognitive decline after stroke, we have developed a framework for future multi-center trials. rCPT: rodent continuous performance task; 5C-SRT: 5-choice serial reaction test; PAL: paired-associated learning; 5C-CPT: 5-choice continuous performance task.

In conclusion, we have demonstrated the inherent challenges in assessing PSCI in pre-clinical stroke using community-established models and cognitive tasks. The NOR task is highly variable, poor performance prior to any surgical interventions leads to a high number of exclusions (which is exacerbated in older mice) and did not detect PSCI in any model at any site. In contrast, the BM is less variable and exclusions due to inability to perform the task prior to stroke are low. Given the promising single-site data from the dMCAO model, the BM warrants testing more broadly. Our findings are informative for the development of multi-site trials to test potential therapeutic interventions and in providing individual labs with important questions to consider when choosing which cognitive tasks may be most suitable in pre-clinical stroke studies. We hope this work will enable the field to move forward towards well-designed pre-clinical studies that are rigorous and reproducible and lead to critically needed therapies to prevent PSCI.

## Acknowledgements

We would like to thank Lizi Hegarty, Michael J.D. Daniels, Tawaun Lucas, Elizabeth Mayne and Li Zhu for their assistance in sample collection. We would also like to thank Amy Bradley for her assistance with network administration, data organization and storage, and coordination between laboratories of the network.

## Author Contribution Statement

GB and KAZ designed and performed experiments, collected and analyzed data, and wrote and edited the manuscript with comments from all authors. DS, JEG, SHL, RC, JBF, DAB, MIC, AGC, CD, DB, and JHF performed experiments, data collection and analysis. AM, JA, MAM, SMA, KPD, MSB and BWM secured funding and established the Stroke-IMPaCT network, designed experiments and edited the manuscript. All authors approved the final manuscript.

## Statements and Declarations

*Ethical Considerations:* All animal protocols were performed in conformity with the Animal Research: Reporting of In Vivo Experiments (ARRIVE) guidelines. Experimental procedures were approved by the appropriate animal care and use committees at each respective institution, and were performed under relevant personal and project licenses.

## Consent for Publication

Not applicable

### Declaration of Conflicting Interest

Dr. Buckwalter has worked as a paid consultant for Roche, Genentech, and EMD Serrano. Otherwise, the author(s) declared no potential conflicts of interest with respect to the research, authorship, and/or publication of this article.

### Funding Statement

The Stroke-IMPaCT network was generously supported by the funding of a Leducq Transatlantic Network of Excellence award to AM, JA, MAM, SMA, KPD, MSB and BWM (19CVD01). This work was also supported by a Knight Brain Resilience award (MSB), the American Heart Association/Allen Frontiers Group Brain Health Award (MSB), Alzheimer’s Association Research Fellowship (KAZ), Grant PID2022-140616OB-I00 funded by MICIU/AEI/10.13039/501100011033 and by ERDF/EU (MAM). GB, DS, BWM are supported by the UK Dementia Research Institute through UK DRI Ltd., principally funded by the UK Medical Research Council. Funding was also provided by the National Institute of Neurological Disorders and Stroke grant RF1NS131110 (KPD) and NS132493 (JA). This work was also supported (AM, DB, CD) by the German Research Foundation (ME1562/4-1; SFB/TRR167: NeuroMac, project B12; Clinical Research Unit KFO 5023: BeCAUSE-Y, project B2). JHF was supported by an Alzheimer’s Research UK senior fellowship (ARUK-SRF2018B-005).

### Data Availability

All data is available upon request from the corresponding authors.

## Notes

### Competing Interest Statement

The authors have declared no competing interest.

