## Supplementary Figures and Tables for "Assessing post-stroke cognition in pre-clinical models: lessons and recommendations from a multi-center study"

**Supplementary Figures & Tables:**

**Supplementary Table 1. Site-specific differences in the novel objection recognition objects.** Adult (12-15 wk old) male C57BL/6J mice were tested on the Novel Object Recognition (NOR) protocol prior to, and 1 and 8 weeks after sham or stroke surgery. Each site had minor variations in objects utilized, and object placement within the testing arena.

| **Site** | **Stroke Model** | **Object Characteristics**  **(LxWxH or DxH cm)** | **Object Position** |
| --- | --- | --- | --- |
| SU | DH | **Baseline**: Lego: 3.2 x 3.2 x 7.7 & 25mL Culture flask: 5.5 x 2.5 x 9.4  **1 wk**: Orange container: 5.3 x 2.5 x 10.5 & 50mL conical: 3.3 x 12  **8 wk**s: green & pink container: 4.0 x 2.5 x 10.5 & White box: 3.2 x 2.9 x 7.5  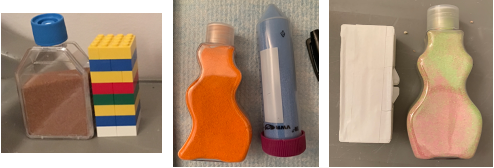 | Center |
| UoA | DH | **Baseline:**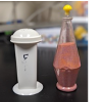 **1 wk:**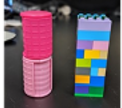 **8 wks:**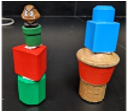 | Top corner |
| UoE | dMCAO | **Baseline**: Lego tower: 3.1 x 3.2 x 7.6 & 50mL falcon tube: 3 x 11.5  **1 wk**: Egg cup: 5.5 x 4.5 x 6.5 & 50mL Culture flask: 4.2 x 4.2 x 10  **8 wks:** Salt shaker:3.8 x 3.8 x 8 & Door knob: 4.6 x 4.6 x 8.2  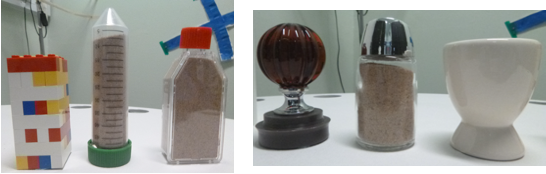 | Top corner |
| WC | tMCAO | **Baseline:** Cylinder: 7.2 x 7.2 x 9.1 & Rectangular box: 3.3 x 8.3 x 9.7  **1 wk:** Pyramid: 5.2 x 8 x 8.5  **8 wks:** Tall square box: 5.7 x 5.9 x 10  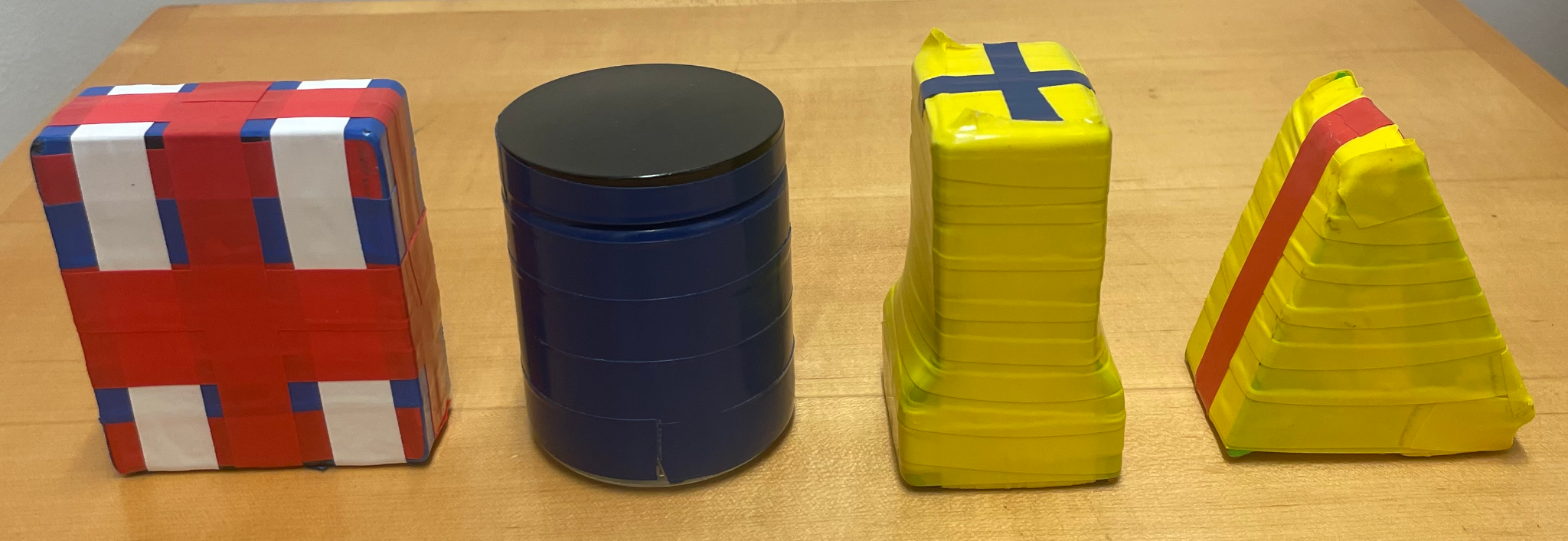 | Center |
| CUB | tMCAO | **Baseline:** Lego tower: 3.2 x 3.2 x 7.7 & 25mL flask: 5.5 x 2.5 x 9.4  **1 wk:**Yellow container: 4.7 x 4.7 x 12 & 50mL falcon tube: 3.3 x 11.5  **8 wks:** Purple container: 9 x 13.5 & white box**:** 8 x 4 x 11  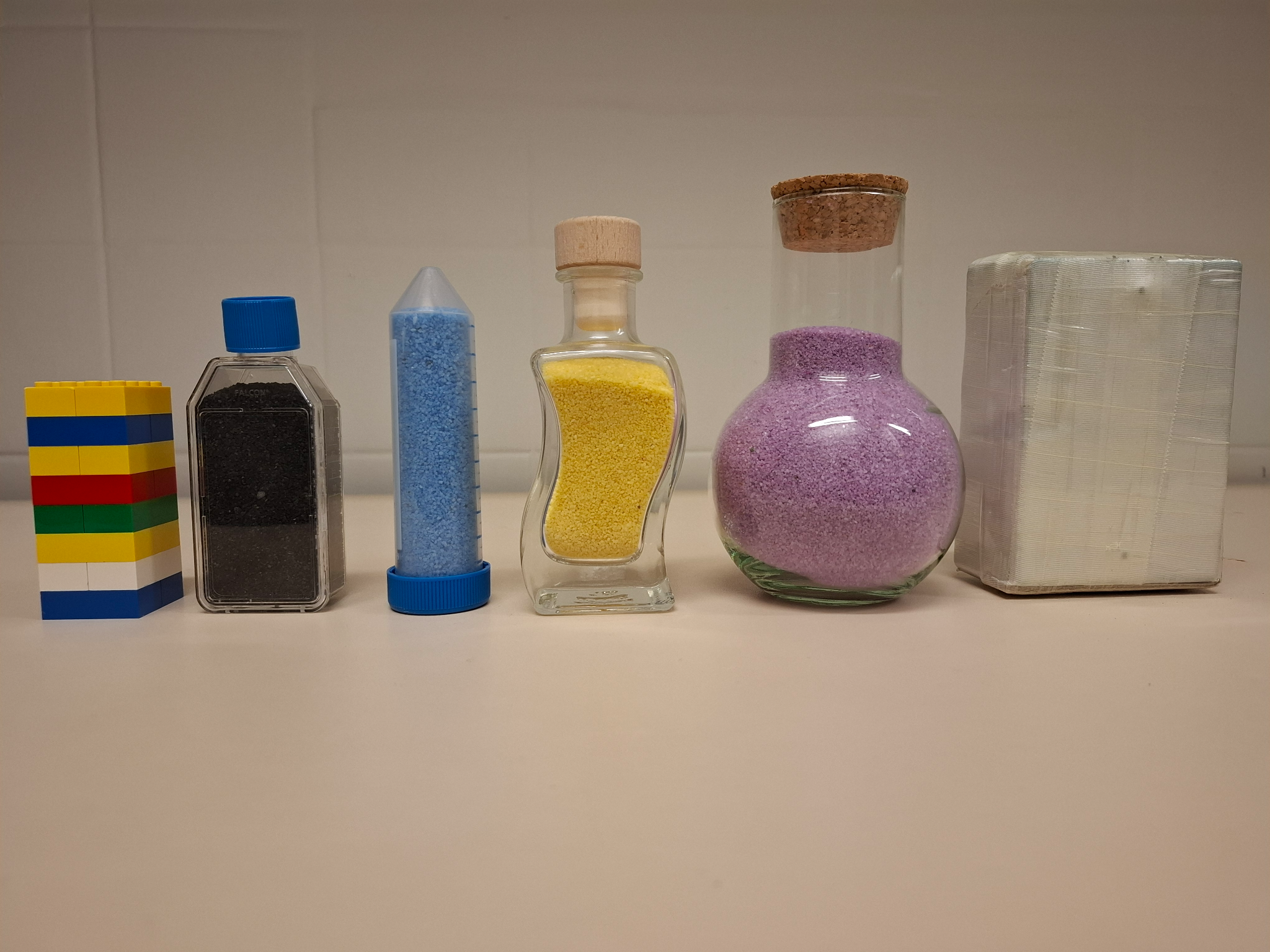 | Center |

**Supplementary Table 2. Potential cognitive behavioral tasks to consider for future pre-clinical studies of post-stroke cognitive impairment.** A ranked list of proposed behavioral tasks, with rationale. Time is time per timepoint.

| **Task** | **Cost** | **Need expert?** | **Time** | **Stroke deficit?** | **Other Comments** |
| --- | --- | --- | --- | --- | --- |
| **Barnes Maze:** Mice are placed on a circular platform with 16 holes around the outside. All are open to the ground except for the escape hole. Mice undergo 4 trials/day over 3-5 days to learn the escape hole location. The primary readout is time to enter the escape hole. Secondary readouts include time to identify the escape hole & time in the target quadrant. | Low | Yes | 3-5d | Yes ^60^ | -sensitive to motor / fatigue deficits  -moderately sensitive to visual acuity change  -may be modified to change difficulty  -ability to test reversal learning |
| **Rodent Continuous Performance Test:** After training on chamber and target identification, 5 images are shown on the screen. Mice must touch the target and keep from touching other images to get rewarded. The primary readout is the average time to touch the correct stimulus. Secondary readouts include time to collect reward, accuracy of selection, and % omitted trials. | High | No | 7d | Yes ^61,62^ | -sensitive to visual acuity changes  -some readouts sensitive to fatigue and anhedonia  -equivalent assessment of processing speed as human CPT |
| **5 Choice- Serial Reaction Test:** After chamber training, mice are rewarded for correctly touching the location of a brief visual stimulus presented in 1/5 random locations. The primary readout is response latency. Secondary readouts include time to collect reward, accuracy, and % omitted trials. | High | No | 7d | No | -some readouts sensitive to fatigue and anhedonia  -can be assessed in humans |
| **Paired Associated Learning**: After chamber training, mice are trained to identify 3 images in a specific location. Each trial includes 2 images, and an 80% chance of presenting a correct image in a correct location, which must be touched by the mouse. The primary readout is time to reach criterion: 80% response accuracy on 2 consecutive days. A secondary readout is accuracy on a pre-defined testing day. | High | No | 7d +  initial time to reach criterion | Yes ^63,64^ | -some readouts sensitive to fatigue and anhedonia  -ability to test reversal learning  -can be assessed in humans |
| **5 Choice-Continuous Processing Test:** After chamber training, mice must identify the location of a brief stimulus presented in 1/5 random locations. Mice must not touch the screen if stimulus is presented in all 5 locations (20% chance). The primary readout is response latency. Secondary readouts include time to collect reward, accuracy, % correctly omitted trials (inhibition), and % incorrectly omitted trials. | High | No | 7d | No | -some readouts sensitive to fatigue and anhedonia  -ability to test reversal learning  -can be assessed in humans |
| **Novel Object Recognition:** In the training phase, mice are habituated with 2 identical objects. During the test phase, mice are exposed to the familiar object and a novel object. The primary readout is % time interacting with each object. Secondary readouts include discrimination index or rears at each object. | Low | Yes | 1-2d | Yes ^65^ | -readout sensitive to motor deficits, fatigue, anxiety, test order  -not robust for use in multiple labs, but works in single labs  -low inter-rater variability for primary read-outs |
| **Fear Conditioning:** Mice are familiarized to a chamber and then conditioned with foot shocks. Retrieval involves testing mice in the same chamber. The primary readout is % time freezing upon retrieval. Secondary readouts include % time grooming. | High | No | 1-2d | Yes ^23^ | -affects performance on subsequent tasks  -task cannot be repeated in the same animals  -no equivalent in people |

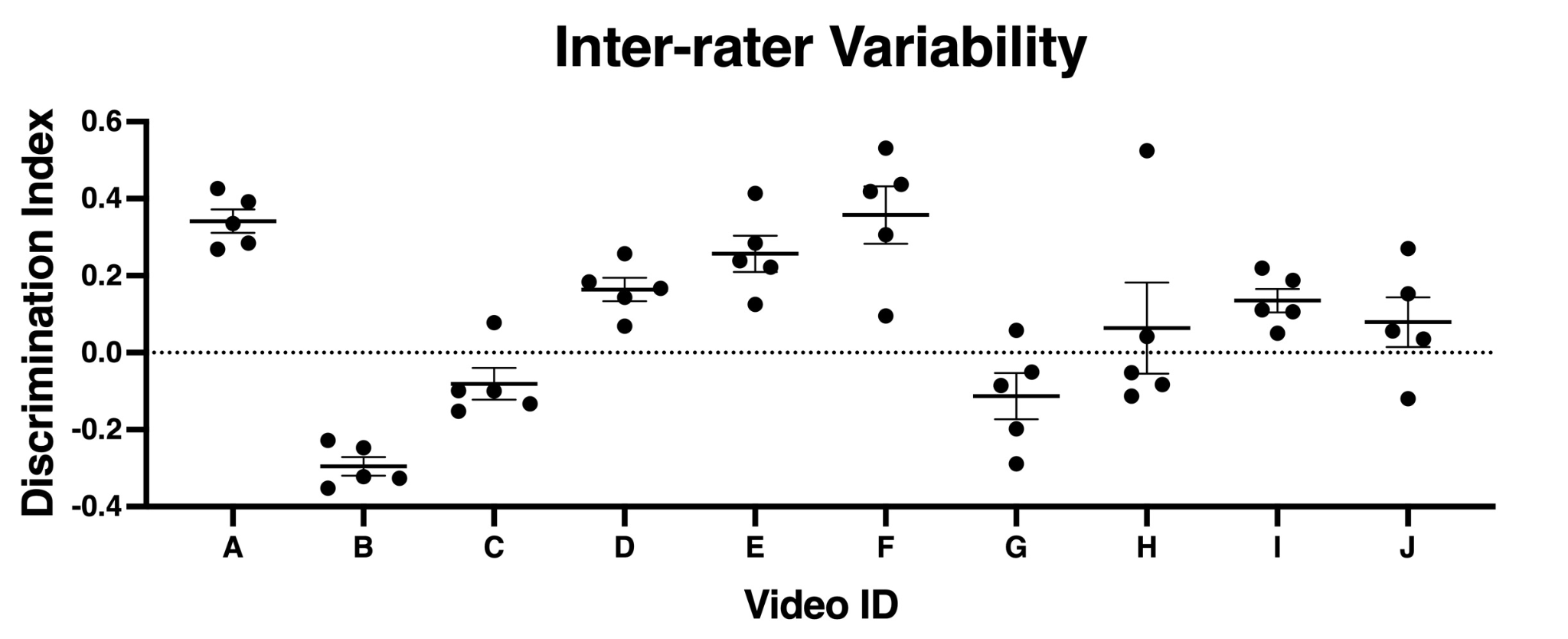

**Supplementary Figure 1. Inter-rater variability**. Discrimination index per video assessed. Each dot represents an assessor. There was one mouse (letters on x axis) per video. Ratings are within 95% CI (correlation coefficient: 0.6991), which met our criteria for rigorous assessment.

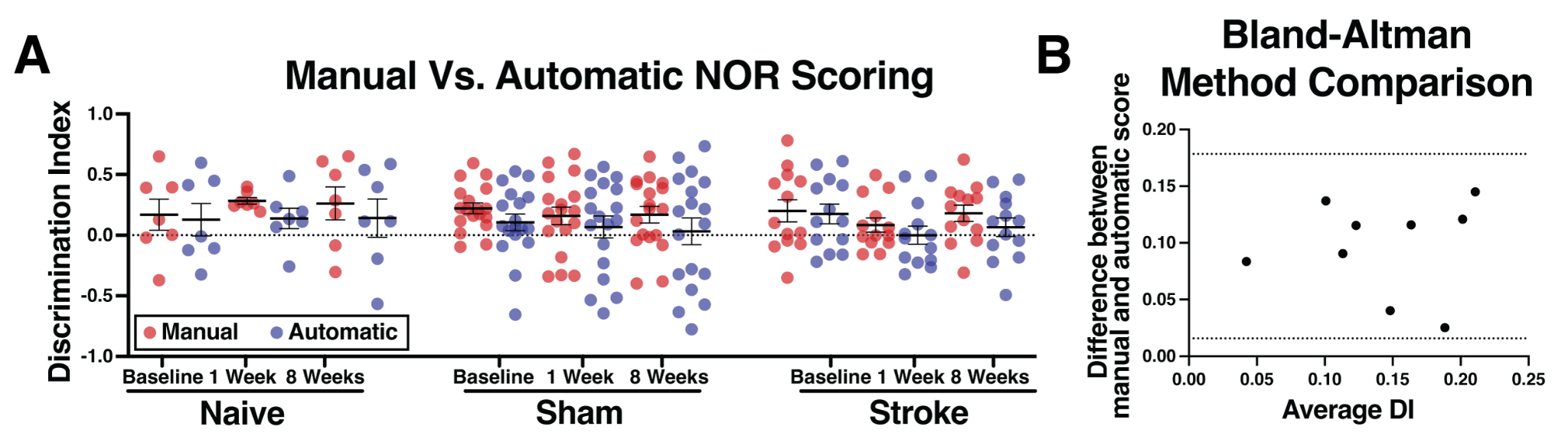

**Supplementary Figure 2.** **(A)** Adult (3 month old) male C57BL/6J mice were tested on the Novel Object Recognition task prior to, and 1 and 8 weeks after DH stroke or sham surgery at the UoA site. Manual scoring was performed by a blinded investigator (red dots) on videos captured during testing, and this was compared to automatic scoring of the same videos using AnyMaze software (blue dots). **(B)** A Bland-Altman method comparison reveals minimal differences in the calculated DI scores between manual and automatic scoring, suggesting that either method can be used for NOR scoring.

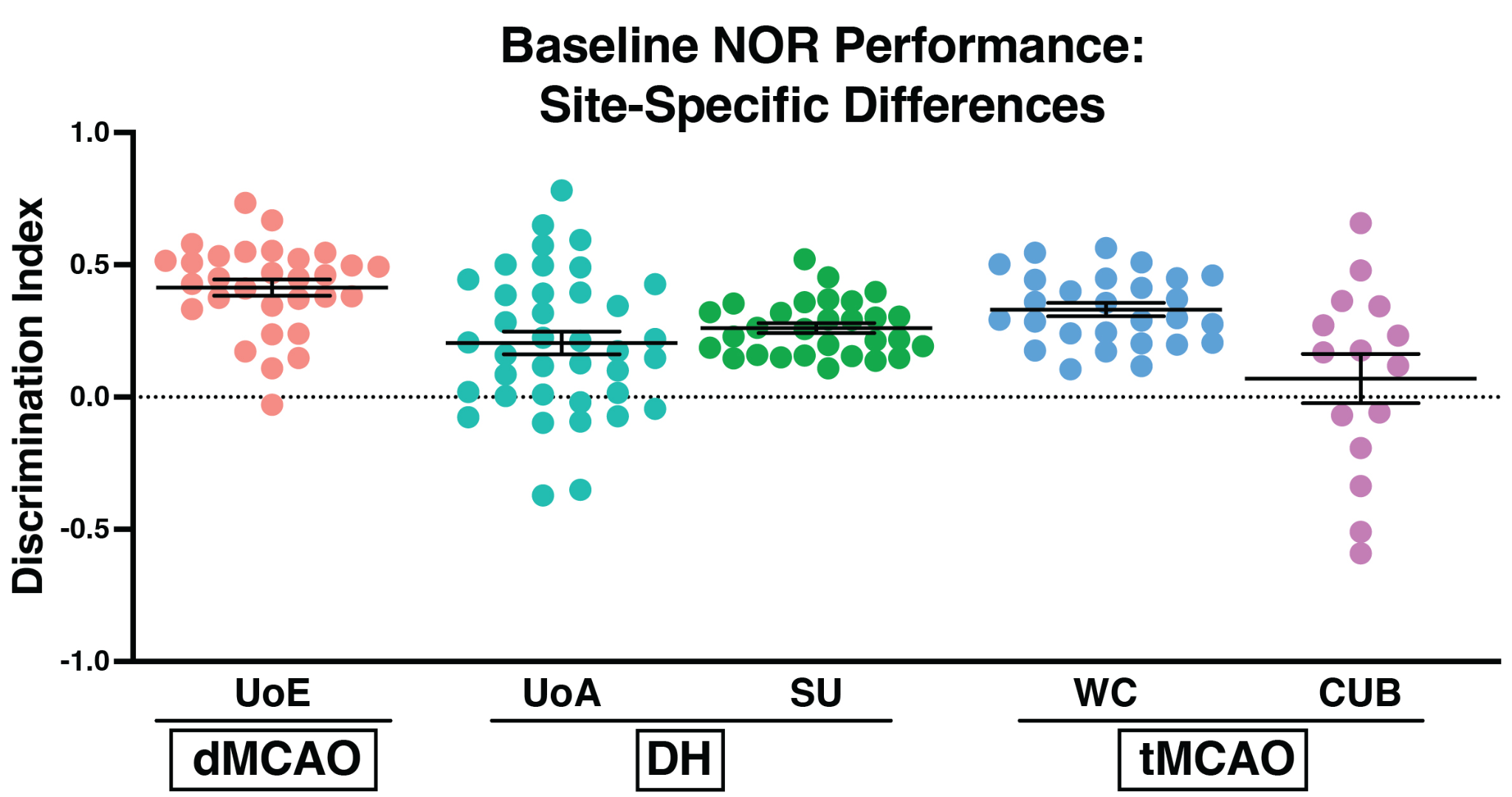

**Supplementary Figure 3. Baseline NOR performance across sites.** Discrimination index (DI) graphed per site prior to any surgical intervention, to establish if each site had a working baseline defined as a DI score of >0.1. Each dot represents an animal.

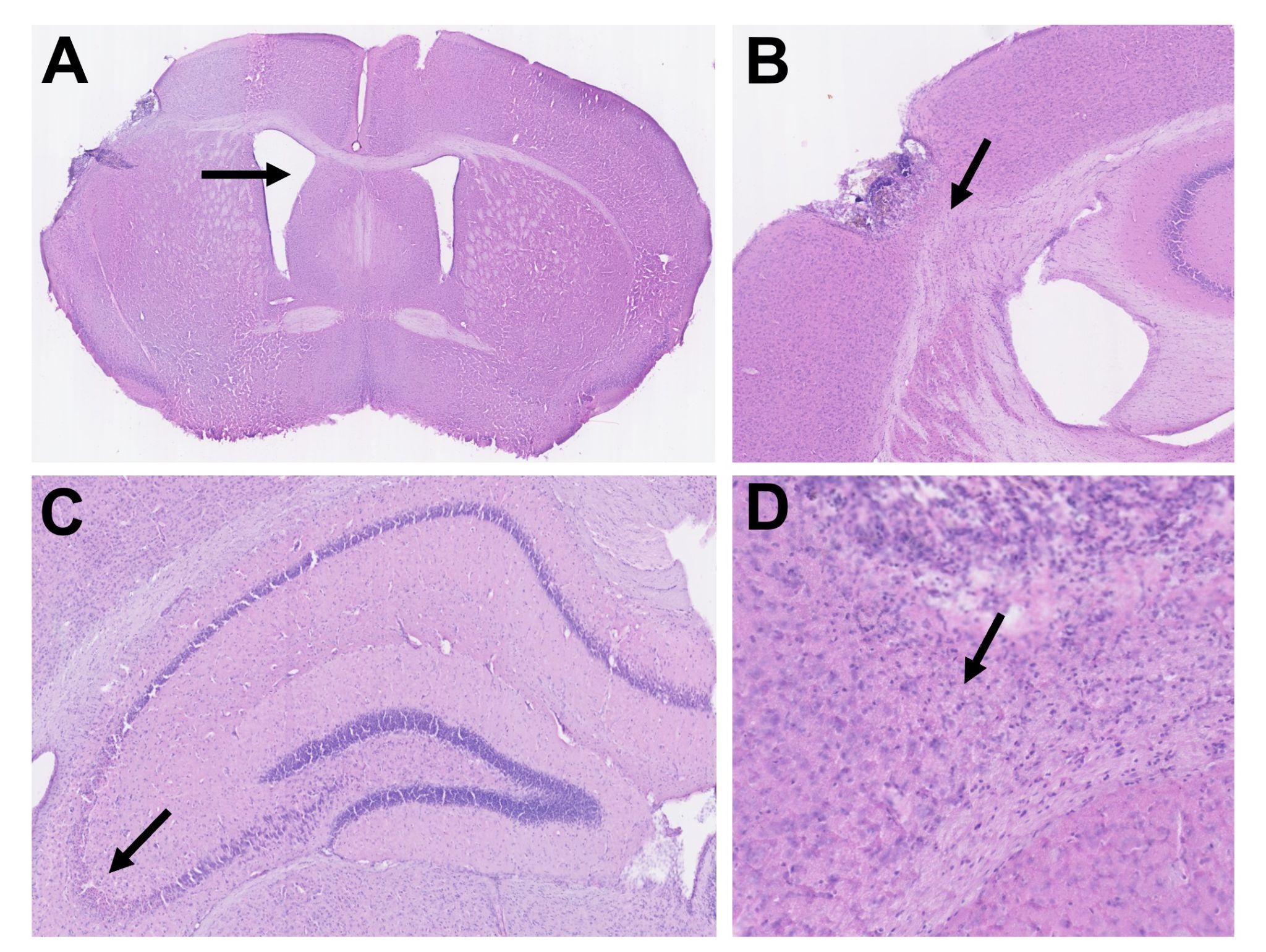
**Supplementary Figure 4. Pathological changes on H&E.** Representative images of pathological features examined by H&E in Figure 2C and 4C. **(A)** Ventricle enlargement. **(B)** Corpus callosum displacement. **(C)** Evidence of neuronal perikaryal shrinkage or damage (i.e hippocampal neuronal death). **(D)** Vacuolation and pallor reflective of pan-necrosis/infarction.

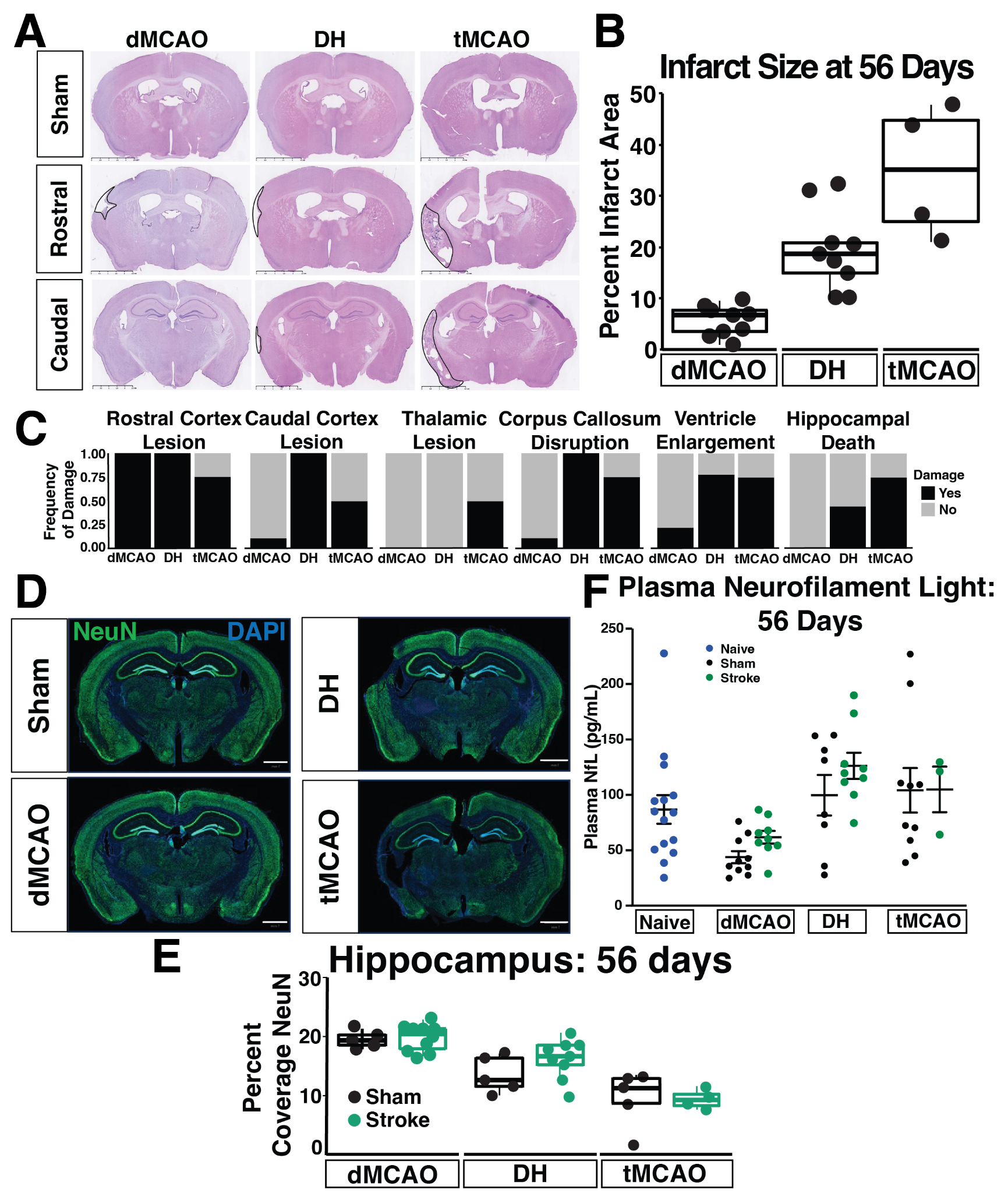

**Supplementary Figure 5. Histopathology assessment at 56 days after stroke in middle-aged mice. (A)** H&E stained representative images from each stroke model (dMCAO, DH, tMCAO) at rostral (Bregma -0.70mm to -0.82mm) and caudal (Bregma -1.70mm to -2.18mm) brain levels. Scale bar=2.5mm. Infarcted tissue is outlined with a solid black line. **(B)** Quantification of infarct size at 56 days post-stroke. **(C)** Assessment of damage frequency at 6 distinct anatomical locations. **(D)** Representative images of whole brain NeuN staining in sham and stroke mice at 56 days post ischemia. Scale bar=1mm **(E)** Quantification of percent NeuN coverage in the hippocampus. (dMCAO, *n*=5-10 mice/group; DH, *n*=5-9 mice/group; tMCAO, *n*=4-5 mice/group). **(F)** Plasma NfL concentration in each stroke model 56 days post ischemia and respective sham controls. Values from naïve mice from all sites (*n*=5 mice/site) are not significantly different and were therefore pooled. Data are presented as mean ± SEM and were analyzed by two-way ANOVA with Tukey’s post hoc test (dMCAO, *n*=9-10 per group; DH, *n*=8-9 per group; tMCAO, *n*=3-10 per group).
