## Supplementary Methods for "Assessing post-stroke cognition in pre-clinical models: lessons and recommendations from a multi-center study"

**Animals**

*Site specific differences in mouse strains, enrichment and diet:*

*Charité Universitätsmedizin Berlin (CUB):* Animals were purchased from Janvier Laboratories (https://www.janvier-labs.com/en/fiche_produit/c57bl-6jrj_mouse/). All animals were housed in open cages in groups of 7-8 mice per cage with a mouse igloo, tunnel, and tissue for enrichment. Mice were fed rat/mouse maintenance diet V1534-000 from www.ssniff.com).

*Weill Cornell Medical College (WC):* Animals were purchased from Jackson Laboratories (https://www.jax.org/strain/000664). All animals were housed in groups of 4-5 mice per cage with nestlets for enrichment. Mice were fed PicoLab Rodent diet 20.

*University of Edinburgh (UoE):* C57BL/6J animals were purchased from Charles River Laboratories. All animals were housed in groups of up to 5 mice per cage with nesting material, tunnel and mouse house for enrichment. Mice were fed Rat and Mouse No.I Maintenance diet from Special Diets Services.

*Centro Nacional de Investigaciones Cardiovasculares (CNIC)*: Animals were purchased from ENVIGO (https://www.envigo.com/model/c57bl-6jolahsd). All animals were housed in groups of up to 5 mice per cage with nesting material for enrichment. Mice were fed LASQCdiet® Rod18-A diet ([www.LASvendi.com](http://www.lasvendi.com)).

*University of Arizona (UoA):* Animals were purchased from Jackson Laboratories (https://www.jax.org/strain/000664). All animals were housed in groups of 4 mice per cage with nestlets for enrichment. Mice were fed NIH-31 Modified Open Formula Mouse/Rat Sterilizable Diet*.*

*Stanford University (SU):* Animals were purchased from Jackson Laboratory, housed in groups of 3-5 mice per cage, with nestlets (https://www.jax.org/strain/000664). Mice were fed Envigo’s Teklad Global 18% Protein Rodent Diet.

**Stroke models**

*For 12-15 week old cohort:* Proximal transient (filament) middle cerebral artery occlusion (tMCAO) was performed at CUB and at WC. Permanent distal middle cerebral artery occlusion (dMCAO) was performed at UoE and at CNIC. dMCAO + Hypoxia (DH) surgery was performed at SU and at UoA.

*For 10-12 month old cohort:* tMCAO was performed at WC. dMCAO was performed at SU and the DH model was performed at UoA. Sham and naïve littermates were used as controls for both age groups, and all mice from all surgical conditions were randomized among experimental cages. Details of stroke and sham equivalent surgery are outlined below. Mice were sacrificed at 3 days or 56 days following stroke or sham surgery, or no surgery (naïve) (**Figure 1A**).

**Surgical procedures by site**

*CUB*: Experimental stroke was induced by proximal middle cerebral artery occlusion using a filament (tMCAO) under 3% isoflurane in a 70/30 mixture of N_2_O/O_2_. Anesthesia was maintained with 1-1.5% isoflurane for the duration of the surgery. Core body temperature was maintained at 37°C with a heating pad. The surgeon made a midline neck incision and exposed the common carotid artery (CCA), external carotid artery (ECA) and internal carotid artery (ICA) on the left side. The vagus nerve was then carefully separated from the blood vessels and the CCA and ECA were ligated with a suture thread. The ICA was temporarily closed with a microvascular clip and an incision was made into the CCA. The surgeon then inserted the filament (Doccol 7-0 fine MCAO suture 701956PK5RE) into the incision and advanced it along the ICA to the origin of the MCA. The filament was secured in the artery with a ligation around the ICA. A drop of 1% bupivacain gel was put into the surgical wound and then temporarily closed. The mouse was placed in a heated cage (30°C) for the 1 h occlusion time. For reperfusion, the mice were re-anesthetized with 2% isoflurane and maintained with 1-1.5% of isoflurane. The surgical wound was reopened and the suture around the ICA opened to retract the filament. Once the filament was retracted, the suture was closed again (CCA, ECA and ICA remain closed) and the wound was sutured. A subcutaneous injection of 0.5 ml 0.9% NaCl was given to prevent dehydration and the mice were allowed to recover in a heated cage (30°C) for 30-60 min before being returned to their home cage. 24 h after surgery, T2-weighted MRI measurements under isoflurane anesthesia were carried out to determine infarct size and location. The surgery for the sham animals was identical to that of the stroke group, except that the filament was immediately withdrawn upon reaching the origin of the MCA.

*WC*: Experimental stroke was induced by proximal middle cerebral artery occlusion (tMCAO) under 3% isoflurane in a 70/30 mixture of N_2_O/O_2_ and maintained with 1.5-2% isoflurane for the duration of surgery. Core body temperature was maintained at 37°C using an electric heating pad. The surgeon made a midline incision behind the ears and the skull was cleaned of connective tissue using a cotton swab. Using instant krazy glue and accelerator, a laser Doppler fiberoptic probe was attached to the skull 2 mm posterior and 5 mm lateral to bregma, and cortical blood flow was recorded. The surgeon then made a midline neck incision. Connective tissue surrounding the CCA, ECA, andICA was cleared and the CCA was isolated from the vagus nerve. Two 6-0 silk sutures (HOSPEQ, Cat #SP114) were tied around the ECA. A third 6-0 silk suture was tied around the CCA, being careful to avoid the vagus nerve, and pressure was applied to reduce blood flow through the CCA, as visualized using laser Doppler flowmetry (LDF). Spring scissors were used to make a small incision in the ECA between the sutures, and 6-0 silicone rubber-coated filament (Doccol, Cat #602112PK10) was inserted and advanced along the ICA to the base of the MCA. The CCA was ligated using a ~1 cm section of PE-50 tubing. The tissue was kept moist using 0.9% NaCl as needed. Following 25 min of ischemia, the PE-50 tubing was removed, and the filament retracted. The sutures around the incision in the ECA remained in place and the suture around the CCA removed to fully restore blood flow. The LDF probe was detached from the skull and both wounds sutured, and 2% lidocaine topical jelly applied. The mice recovered in a heated cage (30°C) for 30-60 min. For seven days after surgery, mice were housed in a 30°C chamber and 0.9% NaCl administered to prevent dehydration as needed. Sham animals underwent the same surgical preparation as stroke mice with the exception of external carotid artery incision and filament insertion.

*UoE*: Experimental stroke was induced by permanent distal middle cerebral artery occlusion (dMCAO) under ~4-5% isoflurane in a mix of 70% N_2_O/30% O_2_ and then maintained by face mask with 1.5-2% isoflurane. Core temperature was maintained at 37 ± 0.5°C with a homeostatic blanket system. The incision site was shaved and cleaned with povidone-iodine antiseptic solution. Lacrllube was applied to the eyes. An incision was made between the left eye and ear and a craniotomy performed, exposing the main trunk and bifurcations of the MCA. The artery was diathermised at 3 locations (at the 3 bifurcations of the MCA or at 3 spaced intervals if no bifurcations are visible). Sham operated mice were not diathermised. Cessation of blood flow was confirmed visually before cutting through the coagulated vessel. Temporal muscle was repositioned, the incision sutured and topical analgesic applied (LMX4 cream). Mice were administered 0.5 ml saline and 0.1 mg/kg buprenorphine subcutaneously, and recovered in a 30°C heat box for 1 h before being returned to home cages. A further subcutaneous injection of 0.1 mg/kg of buprenorphine was administered the morning after surgery.

*CNIC*: dMCAO surgery was performed under 4-5% sevoflurane inhalation, and each mouse received a 100 ml subcutaneous injection of buprenorphine hydrochloride (0.1 mg/kg; Richter Pharma) for pain management. Core body temperature was maintained at 37°C with a surface heating pad throughout the procedure. An incision was then made through the left eye and the ear, and the temporal muscle exposed and retracted. A small craniotomy was then performed over the trunk of the left MCA. Occlusion was performed by ligature of the MCA trunk just before its bifurcation between the frontal and parietal branches with a 9-0 suture (proximal occlusion). Flow disruption was confirmed visually under an operating microscope. The temporal muscle was repositioned and the incision sutured. Sham animals underwent the same procedure, except their MCA was not ligated.

*UoA:* Experimental DH stroke was induced under 3% isoflurane inhalation. Each mouse received a 100 ml subcutaneous injection of buprenorphine hydrochloride (0.1 mg/kg) for pain management (Henry Schein, #2284659). Core body temperature was maintained at 37°C with a surface heating pad throughout the procedure. After shaving the surgical area and swabbing with betadine and sterile saline the surgeon exposed the skull with an incision in the skin and temporalis muscle. Following identification of the right MCA, the surgeon used a microdrill to penetrate the skull and expose the underlying MCA. Then, a small vessel cauterizer (Bovie Medical Corporation) was used to cauterize the MCA. The surgical wound was closed using Surgi-lock 2oc (Meridian Animal Health) and mice were then immediately transferred to a hypoxia chamber (Coy Laboratory Products) containing 9% oxygen and 91% nitrogen for 45 min. After 45 min, mice were returned to their home cages and allowed to recover overnight. At 24 h following surgery, they were administered a 25-50 ml subcutaneous injection of buprenorphine sustained-release at 1 mg/kg for post-operative analgesia (ZooPharm). Sham animals underwent the same procedure, except their MCA was not cauterized.

*SU*: Experimental DH stroke was induced under 1.5-2% Isoflurane in 100% oxygen. Core body temperature was maintained at 37°C with a controlled heat blanket. The incision site was cleaned with chlorhexidine, and then an incision was made along the mouses’ skull, and the temporalis muscle was transected to expose the MCA through the skull. A craniotomy was then drilled, and the artery cauterized. Sham operated mice underwent the same procedure, without artery cauterization. The muscle and skin were replaced, and after closure the animals received cefazolin (25 mg/kg s.c., VWR #89149-888) and buprenorphine SR (1 mg/kg s.c., Zoopharm, Windsor, CO). Animals were allowed to recover for 5 min before spending 60 min in a temperature controlled 37°C chamber in 8% oxygen/92% nitrogen. For dMCAO surgery in older mice, dMCAO or sham surgery was performed as above without exposure to hypoxia, and mice were recovered in a warm cage until awake and mobile.

**Behavioral assessments**

*Novel object recognition (NOR)*

Laboratory specific differences in NOR protocols across sites for 12-15 week old cohorts are reported in Table 1. A unified NOR protocol was used by all sites for testing of the 10-12 month old cohort. Briefly, handling and room habituation procedures were performed at most sites prior to testing (Table 1). The lighting was dimmed in the testing room, and white noise played at some sites (SU). Animals were transferred to the testing room and allowed to acclimate for at least one hour prior to handling, habituation and testing. At some sites (UoE, WC) mice were also habituated to the testing arena prior to testing. Habituation consisted of varying time of exposure to the empty arena as described in Table 1.

The NOR arena was made of acrylic, with objects placed in the center or top corners of the arena (Table 1). The apparatus was positioned in the middle of the room and raised from the floor. The NOR task consisted of two phases: a training phase (Phase I) and testing phase (Phase II). In Phase I, a single animal was placed in the arena with two identical objects. The mouse was allowed to explore freely for 5 min, and the time spent interacting with each object was manually recorded. Interaction was counted when the mouse’s nose was <1 cm (directed within 45 degrees) or touching the object, or its paws were touching the object. Climbing or sitting on an object, or rearing with nose directed >45 degrees away from the object was not considered interaction. After the session, the animal was returned to its home cage. Phase II started 3 h after the completion of Phase I. In this phase, one of the familiar objects was replaced with a novel object. The same animals were reintroduced into the arena individually for 5 min, and their interactions with the objects were recorded with the same criteria above. Following the session, the animal was returned to its home cage. Objects were counterbalanced between animals, to avoid any innate object preference. Upon repeat testing, a new pair of objects was used for each timepoint to avoid potential confounders of long-term object memory. The arena and objects were thoroughly cleaned with 10% ethanol to eliminate residual odors between mice and testing sessions.

During analysis, the following parameters were manually recorded: total exploration time (s) and exploration time (s) with each object. In addition, videos were captured and Phase II object interaction was assessed by at least one other blinded individual from a different site. To formally assess inter-rater variability, 2 videos (1 sham, 1 stroke) were randomly selected from each testing site, and scored by all researchers (6 total scores). One site (UoA) also captured videos and analyzed them via AnyMaze software (version 7, Stoelting Co., Wood Dale, IL, USA) to compare manual to automatic scoring of task performance. To assess performance on the task, a discrimination index (DI) score was calculated with the following formula, (t_novel_-t_familiar_)/(t_novel_+t_familiar_)(tnovel-tfamiliar)/(tnovel+tfamiliar). A score closer to -1 indicates a preference for the familiar object, a score closer to +1 indicates a preference for the novel object, and a score close to 0 demonstrates no object preference.

*Barnes maze (BM) unified protocol*

A modified BM protocol was utilized as previously described ^33^, with minor modifications for 10 month old mice. Prior to the initiation of the experiment, mice were subjected to three days of handling as described above for NOR, and acclimated to the testing room for at least 60 min prior to any handling or testing on a given day.

The BM consisted of a large, circular platform with 16 holes on the outer edge, which was positioned approximately 3 feet off the floor in the center of the room. All of the maze holes were left open to the floor, except for one which contained an escape hole. The escape hole was aligned with one of four distinct visual cues, which were equally spaced around the room. An overhead light, two standing lights, and a fan placed at the edge of the maze were used to motivate animals to find and enter the escape hole.

Mice performed 4 trials per day for 4 days. The escape hole position was fixed for all days of the task, although the starting position of each trial within a day was altered relative to the escape hole position. Each trial had a maximum duration of 90 s. If the mouse failed to find the escape hole within 90 s, the mouse was guided towards the escape hole by tapping its tail and hind legs. Five mice were randomly assigned to each Barnes maze testing group based on cage assignments, and each testing group contained a random mixture of naïve, sham and stroke mice. Each mouse within a testing group was sequentially tested on Trial 1 before moving on to Trial 2. All four trials were completed for Testing Group 1 before proceeding to Testing Group 2, and this sequence was repeated for each testing group until all testing groups completed 4 trials each day. The order of testing, groups and start time of testing were maintained on each day to reduce circadian variability. All equipment was wiped down with 10% ethanol before introducing the next animal, and 70% ethanol at the end of the testing day. The time to identify the escape hole (primary latency; s) and the time to enter the escape hole (escape latency) were manually recorded.
